# Gene knockdown via electroporation of short hairpin RNAs in embryos of the marine hydroid *Hydractinia symbiolongicarpus*

**DOI:** 10.1101/2020.04.07.030296

**Authors:** Gonzalo Quiroga-Artigas, Alexandrea Duscher, Katelyn Lundquist, Justin Waletich, Christine E. Schnitzler

## Abstract

Performing gene function analyses in a broad range of research organisms is crucial for understanding the biological functions of genes and their evolution. Recent studies have shown that short hairpin RNAs (shRNAs) can induce gene-specific knockdowns in two cnidarian species. We have developed a detailed, straightforward, and scalable method to deliver shRNAs into fertilized eggs of the hydrozoan cnidarian *Hydractinia symbiolongicarpus* via electroporation, yielding gene-targeted knockdowns that can be assessed throughout embryogenesis, larval settlement, and metamorphosis. Our electroporation protocol allows for the transfection of shRNAs into hundreds of fertilized *H*.*symbiolongicarpus* eggs simultaneously with minimal embryo death and no long-term harmful consequences on the developing animals. We show RT-qPCR and detailed phenotypic evidence of our method successfully inducing significant knockdowns of an exogenous gene (*eGFP*) and an endogenous gene (*Nanos2*). We also provide visual confirmation of successful shRNA transfection inside embryos through electroporation. This is the first time that electroporation as a delivery system has been developed for *Hydractinia*. Our detailed protocol for electroporation of shRNAs in *H. symbiolongicarpus* embryos constitutes an important experimental resource for the hydrozoan community while also serving as a successful model for the development of similar methods for interrogating gene function in other marine invertebrates.

---

Hydrozoans are members of the phylum Cnidaria, a group that holds a phylogenetic position as sister to all bilaterian animals^1^ (Fig. **1A**), and thus can provide insights into the origins of key bilaterian features. Research involving hydrozoans has greatly contributed to our understanding of crucial cellular and developmental processes, as well as their evolution^2^. Within the Hydrozoa, members of the genus *Hydractinia* have been used as experimental research organisms for more than a century^3–5^. *Hydractinia* is a dioecious, marine, colonial hydroid that is well-suited for lab culturing and rearing. Its small size and transparency make it tractable and convenient for manipulation and microscopic imaging. Moreover, its embryonic development has been well-described^6^, allowing for targeted developmental studies. A fundamental characteristic of *Hydractinia* is that it maintains a population of stem cells called interstitial cells (or ‘i-cells’) that provides progenitors to both somatic and germ cell lineages in a continuous manner throughout its lifetime, allowing for remarkable regenerative capabilities and longevity^3,4,7^. The availability of a sequenced genome, and the range of functional genomic tools currently available make *Hydractinia* a rapidly maturing cnidarian system that is allowing researchers to explore a wide array of biological topics, ranging from stem cells and regeneration to developmental biology and self-recognition (allorecognition)^3–5,8^. In this study, we focused on a new method to silence genes in the species *Hydractinia symbiolongicarpus* (Fig. **1B**), whose life cycle is shown in **Figure 1C**. The relatively short and accessible life cycle of *H. symbiolongicarpus* allows for experimental manipulation of all life stages.

**Figure 1.**
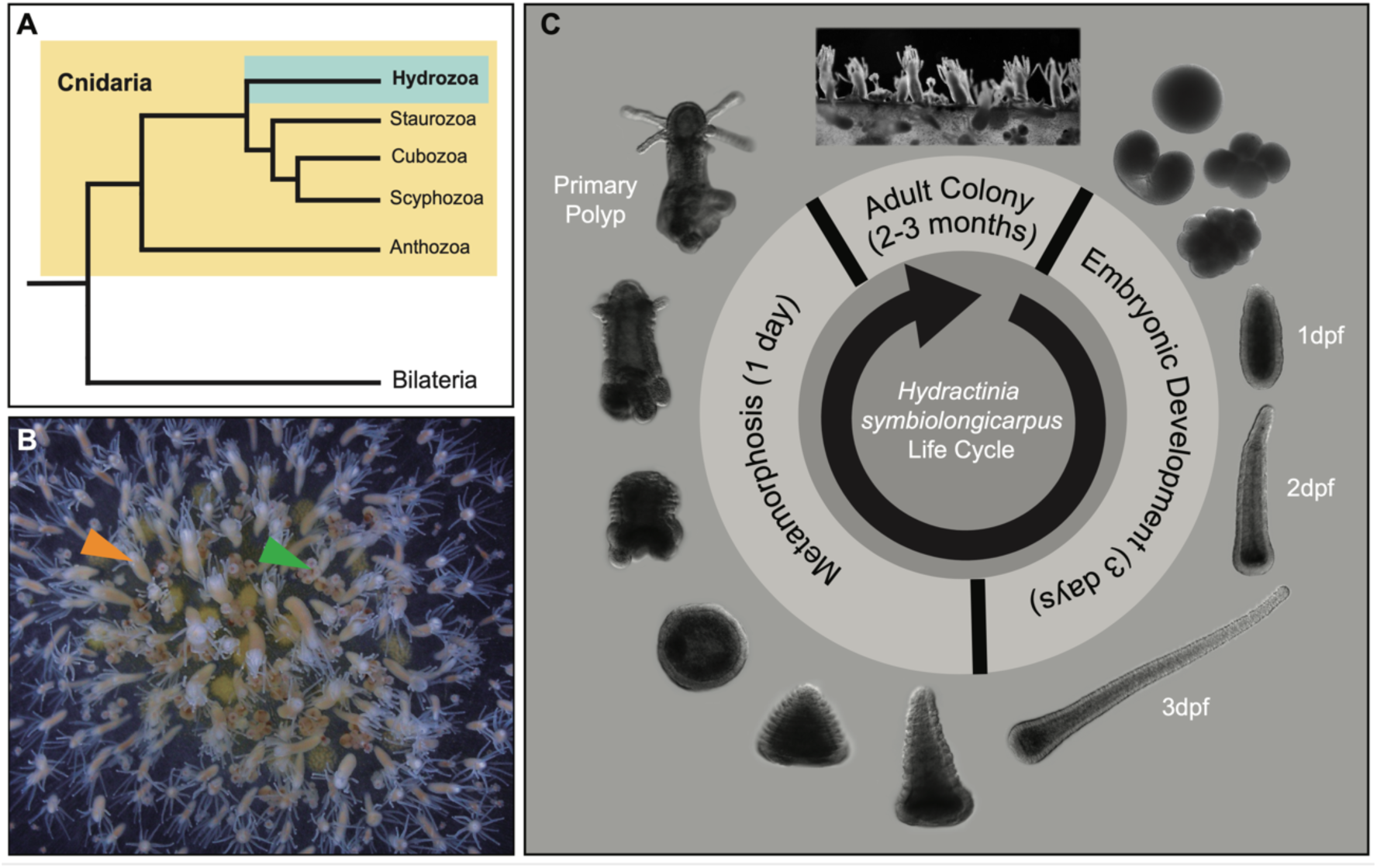
The hydrozoan cnidarian *Hydractinia symbiolongicarpus*. (**A**) Cladogram depicting the phylogenetic position of Cnidaria as sister group of Bilateria, as well as the phylogenic relationships between Hydrozoa and other cnidarian groups, based on Kayal *et al*.^44^. (**B**) Photo of a female adult colony of *H. symbiolongicarpus*. Orange arrowhead points to a feeding polyp and green arrowhead points to a reproductive polyp. (**C**) *H. symbiolongicarpus* life cycle. Images show key stages of embryonic development as well as stages of metamorphosis from a larva to a primary polyp and ultimately the adult colony. Mature female and male colonies spawn hundreds to thousands of gametes daily following a light cue. Spawned eggs are fertilized in the water column, and proceed through embryonic development^6^. About three days post-fertilization (3 dpf), fully-developed larvae are competent to receive a natural or artificial stimulus that will induce settlement and metamorphosis^45,46^. Metamorphosis is completed within 24 hours, and this process transforms a mouthless larva into a feeding primary polyp. The animal then expands clonally by stolonal elongation and asexual budding of new polyps. The polyps are interconnected by a stolonal mat which holds gastrovascular canals, enabling food distribution and transfer of stem cells throughout the growing colony^47^. Two to three months after metamorphosis, reproductive polyps start to bud, allowing the colony to reproduce sexually.

The ability to experimentally alter a gene’s function is crucial for understanding its biological role. In the last decade, a variety of techniques to study gene function have become available for *Hydractinia*. Most recently, genetic engineering (through both gene knockouts^9^ and knockins^10^) has been achieved in *Hydractinia* by means of CRISPR/Cas9 embryo microinjection. Prior to the advent of these gene editing techniques, antisense RNA-mediated gene silencing to induce gene-specific knockdowns had been frequently employed in *Hydractinia*, including morpholino microinjection in embryos^11^ and double-stranded RNA (dsRNA) soaking in embryos and polyps^7,12–15^.

dsRNAs exploit the endogenous RNA interference (RNAi) machinery of the cells and have been used as molecular tools to transiently lower the transcript levels of a particular target gene (i.e., knockdown) for decades^16^. RNAi knockdown offers some distinct advantages over gene editing knockout techniques – in particular, the avoidance of loss-of-function lethality caused by the complete disruption of certain genes, since RNAi transiently reduces but does not entirely remove gene activity^17^.

Short hairpin RNAs (shRNAs) are small, synthetic dsRNA molecules connected by a hairpin loop that can be used instead of longer dsRNAs to knock down target genes via RNAi^17^. shRNAs are processed similarly to precursor microRNAs (pre-miRNAs) through the endogenous RNAi pathway of transfected cells. The enzyme Dicer cleaves the loop and converts shRNAs into siRNAs (small interference RNAs). The resulting siRNAs are unwound into passenger (forward) strands, which are degraded, and guide (reverse) strands, which are incorporated into the RNA-induced silencing complex (RISC). The guide strand-RISC complex binds to the targeted mRNAs, and cleaves them or inhibits their translation^17,18^. shRNAs have been widely used to induce gene knockdowns in mammalian cell culture^19^ and model systems such as the fruit fly^20^ and zebrafish^21^. This knockdown approach has recently been used to target a small number of genes in two cnidarian species^22–24^. In contrast to bilaterian systems, it has been shown that cnidarian miRNAs require perfect to nearly perfect complementarity between the miRNA guide strands and their target mRNAs to function properly^22,25^, commonly leading to slicing of the target mRNAs^25^. Two studies using the anthozoan cnidarian *Nematostella vectensis* capitalized on this specificity of the cnidarian miRNA pathway and demonstrated gene-specific knockdowns by delivering *in vitro-*synthesized shRNAs inside unfertilized eggs either via microinjection^22^ or electroporation^23^. In a more recent study using *H. symbiolongicarpus*, shRNAs microinjected in fertilized eggs successfully yielded targeted gene knockdown^24^, making shRNAs a promising tool to silence genes in *Hydractinia*.

One of the factors currently limiting the wider application of shRNAs as a gene silencing strategy in *Hydractinia* and other hydrozoan embryos is the lack of a method for easy, fast, scalable, and efficient shRNA delivery. Microinjection as a delivery method is limited by the number of eggs that can be injected and the number of conditions that can be examined in the course of a single experiment. This is particularly problematic when working with fertilized eggs, which begin to cleave in under an hour after fertilization occurs^6^. Delivery via soaking requires constant incubation with relatively high concentrations of the antisense molecules, an approach that can be expensive and may not be consistently effective due to potential cell permeability changes throughout an experiment. As an alternative to these delivery methods, electroporation has been widely used by developmental biologists to transfect embryos from different phyla with the desired biomolecules^26^, most recently to transfect *Nematostella* unfertilized eggs with shRNAs^23^. Electroporation is a physical transfection method that temporarily increases cells membrane’s permeability when submitted to electric field pulses^27^. The major advantage of this delivery method is that it is very fast, allowing for the transfection of many hundreds of eggs at one time, as well as being able to examine a larger number of experimental conditions per assay. One of the challenges of electroporation is balancing efficient transfection and high survivorship, which involves finding conditions that allow the embryos to survive the transfection process. The specific electroporation parameters (voltage, number of pulses, and pulse length), therefore, must be optimized for each species, life cycle stage, and biomolecule type.

Here, we describe a detailed, optimized, and accessible method for simultaneously transfecting hundreds of embryos with shRNAs via electroporation in *H. symbiolongicarpus* – the first report of the successful application of this methodology in embryos from any hydrozoan cnidarian species. We demonstrate that this procedure can successfully induce the knockdown of both exogenous and endogenous targeted genes throughout embryonic development, settlement, and metamorphosis. We also provide visual confirmation of the successful delivery of shRNAs inside embryonic cells upon electroporation and a 28-day time series experiment showing that there are no long-term harmful consequences on the shRNA-transfected animals. This new method will enable researchers interested in different facets of hydrozoan biology to study the function of targeted genes in a simple and scalable manner, and represents a significant experimental advance that will be of benefit to the broader community of researchers working with a variety of cnidarian and other marine invertebrate species.

## Methods

### Animal care and breeding

*H. symbiolongicarpus* adult colonies were maintained at the University of Florida’s Whitney Laboratory for Marine Bioscience. Colonies were grown on glass microscope slides and cultured in 38-liter tanks filled with artificial seawater (Instant Ocean - Reef Crystals) at 30 ppt and kept at 18–20°C under a 10h/14h light/dark regime. Animals were fed three times a week with 3-day-old brine shrimp nauplii cultured at 25°C (Premium Grade, Brine Shrimp Direct), which had been enriched with Shellfish Diet (Reed Mariculture) one day after hatching, and twice a week with an oyster puree made from freshly caught, shucked, and blended oysters (stored at -20°C).

Spawning of male and female gametes was induced by light stimulation. Sperm and eggs were mixed together to allow fertilization (detailed procedure in Supp. File S1). All embryos were allowed to develop into planula larvae in 100 mm×15 mm glass petri dishes. When necessary, 72 hpf (hours post-fertilization) larvae were induced to metamorphose by incubating them in 116mM CsCl solution in MFSW (Millipore-filtered seawater) for 3h, as previously described^8^, followed by two washes in MFSW, and finally transferring them with gelatin-coated glass pipettes onto 75×25mm glass microscope slides for settlement. Primary polyps that were kept for longer than two days post-metamorphosis were mouth-fed smashed brine shrimp every other day.

### shRNA design and synthesis

shRNA design was generally performed as previously described for the cnidarian *Nematostella vectensis*^22,23^. A detailed protocol of our design strategy with examples can be found in Supp. File S2. The DNA templates for *in vitro* transcription of shRNAs are composed by the T7 RNA polymerase promoter at the 5’ end, followed by a forward and reverse strand targeting the gene of interest, which are in turn connected by the TTCAAGAGA hairpin loop linker, and two additional thymidines at the 3’ end to mimic the endogenous pre-miRNA structure. Forward and reverse oligos of 66 bases in length corresponding to the DNA templates for *in vitro* transcription of shRNAs (Supp. Table S1) were ordered from ThermoFisher Scientific, resuspended in nuclease-free H_2_O to a concentration of 100μMol and stored at -20°C.

shRNA synthesis was carried out as previously described^23^ with minor modifications. Briefly, dsDNA templates for *in vitro* transcription of shRNAs were constructed by mixing 2μl of each of the two 66-base oligos (forward and reverse;100μMol each) with 16μl of nuclease-free H_2_O, followed by a 2min incubation at 98°C and a 10min incubation at RT (room temperature). *In vitro* transcription of shRNAs was performed using the AmpliScribe™ T7-Flash™ Transcription Kit (Lucigen). The transcription reaction was allowed to continue at 37°C for 5-7h. An incubation with DNase at 37™C for 30min was included at this point to eliminate the DNA template. Newly synthesized shRNAs were then purified using the Direct-zol™ RNA MiniPrep Kit (Zymo Research), eluted in 35μl of nuclease-free H_2_O and quantified with a NanoDrop 2000 spectrophotometer (ThermoFisher Scientific). Normally, two synthesis reactions per shRNA were performed in parallel and pooled through the same purification column to attain higher concentrations of shRNAs. Purified shRNAs were stored at -80°C for up to four months.

### Electroporation procedure

A detailed, step-by-step protocol can be found in Supp. File S1. Briefly, 300-800 fertilized eggs (also referred to as one-cell stage embryos) were transferred to a well in a glass depression slide and seawater was removed and replaced with 100μl of the electroporation mixture (15% Ficoll in MFSW containing shRNAs; Dextran; Or nuclease-free H_2_O at given experimental concentrations). Ficoll is a large synthetic polysaccharide that has been previously shown to make embryos float, allowing a more homogeneous voltage delivery, when mixed with seawater at 15% concentration^23^. Next, embryos were transferred into an electroporation cuvette (2mm gap) and placed inside the safety stand connected to the ECM 830 square wave system (BTX). Embryos were then subjected to experimental electroporation parameters. When more than one pulse was delivered, the time span between pulses was always kept at 0.5 seconds. After a few seconds, electroporated embryos were gently transferred to a 100 mm×15 mm glass petri dish filled with MFSW and left undisturbed to recover and develop for several hours (Fig. **2**). Dead and non-developing embryos were removed once a day, including on the day of electroporation, to allow for better development of the survivors. MFSW was also changed on a daily basis. All embryos were kept at ∼18°C throughout experimental procedures and subsequent development. Non-electroporated (NE) controls were prepared several times by soaking the embryos in the *eGFP* shRNA mixture at 300 ng/μl each for the same length of time as the electroporation procedure (∼3 minutes). In all cases, we observed that the NE control yielded no obvious fluorescence reduction or *eGFP* knockdown. Therefore, to avoid wasting shRNAs, we used the same proportions of nuclease-free H2O instead of shRNAs in the NE controls for all subsequent experiments with *eGFP* or *Nanos2*.

### Light and fluorescence imaging

Images taken for survivorship and eGFP (enhanced green fluorescent protein) fluorescence analyses as well as for Supp. Fig. S1 and S2 were taken with a digital camera (Canon DS126201) attached to a stereo microscope (Zeiss, Discovery.V8). Light and fluorescence images shown in Fig. 1, 3, 4 and 6 were taken with a Rolera EM-C^2^ high-speed camera (QImaging) attached to a fluorescence microscope (Zeiss, Imager.M2). Identical scanning parameters (i.e., magnification and exposure time) were used for all conditions for each independent experiment.

### eGFP fluorescence and survivorship analyses

All image processing was done using ImageJ software^28^. To count eGFP^+^ larvae, the image background was enhanced identically for all images, and individuals were highlighted using identical thresholding for all conditions, separated using a watershed filter setting, and counted with the ‘Analyze Particles’ tool (Supp. Fig. S2). To assess survivorship, the same procedure was carried out, without background enhancement, for all life cycle stages assessed (Supp. Table S2).

### RT-qPCR

To quantify shRNA knockdown of the targeted genes, total RNA was extracted with the RNAqueous(tm)-Micro Total RNA Isolation Kit (Ambion) at different timepoints from larvae or primary polyps, depending on the experiment. For each condition from every experiment, between 250 and 700 individuals were collected for RNA extraction.

Samples were then treated with DNase according to manufacturer’s recommendations, and the RNA was reverse transcribed with random primers using the High Capacity cDNA Reverse Transcription kit (Applied Biosystems). qPCR analyses were performed using the PowerUp SYBR Green Master Mix (Applied Biosystems) and the LightCycler® 480 instrument (Roche). For each gene and condition, expression levels were derived from three amplification reactions, and normalized to housekeeping gene expression, either to *18S* rRNA (*eGFP* experiments) or to *Eef1alpha* (*Nanos2* experiments). The delta-delta-ct method was used for quantifications of transcript levels from experimental conditions relative to scrambled shRNA or NE (non-electroporated) controls. These relative expression levels are shown as absolute units (A.U.) in all figures. At least three independent biological replicates were performed for each experiment. Primer sequences can be found in Supp. Table S1.

### Dig-shRNA synthesis and tyramide signal amplification

Digoxigenin-labeled shRNAs (Dig-shRNA) were synthesized as described above, using a nonspecific scrambled sequence as dsDNA template, and adding 0.5μl UTP and 1.5μl Digoxigenin-11-UTP (Roche) in the synthesis reaction.

Embryos were electroporated as discussed above with Dig-shRNA at 900 ng/μl and the appropriate negative controls were included (see Fig. **5**). Four hours after fertilization, corresponding to the 32-64 cell stage embryonic stage, embryos were fixed overnight at 4°C in 4%PFA-MFSW (Millipore-Filtered Seawater buffer containing 4% paraformaldehyde). Then, samples were washed five times in PBS containing 0.1% Tween20 (PTw), dehydrated and permeabilized by increasing concentrations of methanol in PTw (25%, 50%, 75%, 100%), and stored at -20°C for at least 24h.

Samples were rehydrated by decreasing concentrations of methanol in PTw (75%, 50%, 25%), washed three times in PTw, followed by two washes in PBS-0.02% Triton X-100, then one time in PBS-0.2% Triton X-100 for 20min, and again two times in PBS-0.02% Triton X-100. They were then blocked in PBS with 3% BSA for 3h at RT and incubated overnight at 4°C with a peroxidase-labeled anti-DIG antibody (Roche) diluted 1:1500 in PBS with 3% BSA. Samples were then washed six to eight times for 15min in PBS-0.02% Triton X-100 and incubated for 30min at RT in Tyramide buffer (NaCl at 116.88 mg/ml and Boric Acid at 6.18 mg/ml in nuclease-free H_2_O; pH = 8.5). Selected samples were incubated for 30min at RT in development solution (Tyramide buffer, 0.0015% H_2_O_2_, NHS-Rhodamine at 4μg/ml) to perform the tyramide signal amplification (TSA). All samples were washed six times in PTw and the last wash was left overnight at 4°C. Nuclei were stained using Hoechst dye 33342 and samples were mounted in Fluoromount® (Sigma-Aldrich).

### Nematocyst staining

Staining of nematocysts was performed as previously described^29^. Larvae were anesthetized in 4% MgCl_2_ in 50% MFSW / 50% H_2_O for 30min before fixation. Samples were fixed in nematocyst fix (10mM EDTA, 4% PFA in PTw) at RT for 1h, then washed three times in PTw containing 10mM EDTA. Next, samples were incubated with DAPI at 1.43μM in PBS1x overnight at 4™C. This was followed by five to six washes in PTw with 10mM EDTA, and samples were mounted in Fluoromount® prior to imaging.

### Immunofluorescence and Phalloidin staining

Larvae were anesthetized in 4% MgCl_2_ in 50% MFSW / 50% H_2_O for 30min prior to fixation. For immunofluorescence, samples were fixed at RT for 2h in HEM buffer (0.1 M HEPES pH 6.9, 50 mM EGTA, 10 mM MgSO_4_) containing 0.02% Triton X-100% and 4% paraformaldehyde in MFSW, and then washed four times in PBS-0.3% Triton X-100. The last wash was left rocking overnight at 4™C. Samples were then blocked in 3%BSA / 5% goat serum in PBS-0.3% Triton X-100 for 3h at RT and then incubated in primary antibody (anti-GLWamide^30^) diluted to 1:200 in blocking solution overnight at 4™C. This was followed by four washes in PBS-0.3% Triton X-100 before samples were blocked again in blocking solution for 1h at RT, and then incubated with secondary antibody (goat anti-rabbit 556; Invitrogen) at a concentration of 1:500 in blocking solution for 1h at RT. Animals were then washed four times for 15min in PBS-0.3% Triton X-100 prior to nuclei staining using Hoechst dye 33342. Samples were mounted in Fluoromount® prior to imaging.

For Supp. Videos S1 and S2, embryos were electroporated with Dextran (Alexa Fluor 555; Invitrogen) at 1mg/ml in Ficoll 15% MFSW and fixed 3 days post-electroporation in 4%PFA in PTw for 1h at RT. Samples were washed in PBS-0.2% Triton X-100 for 15min followed by several washes in PTw. To visualize F-actin, samples were incubated for 2h in Phalloidin (Invitrogen, Cat. A12379) diluted 1:100 in PTw. Samples were then washed four times in PTw, followed by nuclei staining using Hoechst dye 33342. Samples were mounted in Fluoromount® prior to imaging.

### Confocal microscopy and cell counting

Images were acquired using a Zeiss LSM 710 confocal microscope. The same scanning parameters (i.e., magnification, laser intensity and gain) were used for all conditions of each independent experiment. All supplementary videos as well as maximum intensity projections of z-stacks were prepared using ImageJ software^28^.

For DAPI-stained nematocysts, confocal z-stacks of ∼10μm focused on the larval surface were projected. Nematocysts were highlighted using custom thresholding, separated using a watershed filter and counted with the ‘Analyze Particles’ tool from ImageJ. For GLWamide^+^ neurons, confocal z-stacks of ∼45μm focused on the larval aboral region were projected and neuronal bodies were counted manually using the ImageJ ‘cell counting’ plugin.

### Graphs and statistical analyses

Box plots for Fig. **7** and Supp. Fig. S3 were prepared using BoxPlotR^31^. All other graphs were prepared in Excel and assembled using Adobe Illustrator.

To assess RT-qPCR statistical significance, we used the delta-delta-Ct values to perform Shapiro-Wilk normality tests and two-tailed Student’s T-Tests. For statistics related to counts of nematocysts and GLWamide^+^ neurons (Fig. **7**), Kolmogorov-Smirnov normality tests were used, and two-tailed Student’s T-Tests were performed. For tentacle number comparisons (Supp. Fig. S3), the nonparametric Mann-Whitney U test was chosen since the results did not follow a normal distribution according to the Kolmogorov-Smirnov test, and significance between sample distributions could be appropriately assessed by data transformation into ranks. All statistical tests were performed at http://www.socscistatistics.com.

## Results

### Initial electroporation trials with Dextran

We first tested the efficiency of Dextran transfection into *H. symbiolongicarpus* one-cell stage embryos using different electroporation conditions. Dextrans are polysaccharides that can be synthetically labelled, and they have previously been used as long-term tracers in cnidarians^22,32^. The Dextran (Alexa Fluor 555) used in our experiments fluoresces when excited with green light and has a similar molecular mass to shRNAs (10,000 Da and ∼15,000 Da, respectively). We reasoned that electroporation trials with Dextran would help to visually demonstrate whether small molecules can be successfully transfected into *H. symbiolongicarpus* embryos and to test conditions to maximize the survivorship of embryos and overall transfection efficiency.

Our initial electroporation trials were performed with unfertilized eggs that were subsequently fertilized, but we found that fertilizing after electroporation led to almost no healthy embryos. We performed all subsequent trials with one-cell stage fertilized embryos, which resulted in high percentages of healthy, developing embryos. We tested six different electroporation conditions (conditions 1-6), modifying the voltage (volts – V), the number of pulses, and/or the pulse length (milliseconds – ms) in each trial (Supp. Fig. S1). Electroporation trials were carried out as described in **Figure 2**, but instead of shRNAs, Dextran was diluted in 15% Ficoll MFSW at a concentration of 1mg/ml. We defined successful electroporation conditions as those that visually yielded high levels of Dextran fluorescence as well as high embryo survival rates at 1 day post-fertilization (dpf). A non-electroporated (NE) control was added, where samples were soaked in Dextran diluted in 15% Ficoll MFSW at the same concentration for ∼3 min, the approximate length of an electroporation procedure, to allow appropriate comparisons. We found that the 1 dpf larvae in the NE controls were almost non-fluorescent and the survival rate was near 100%, as expected (Supp. Fig. S1). Of the six electroporation conditions tested, conditions 1-3 showed the highest levels of fluorescence in 1 dpf larvae. Condition 3 (25 V, 1 pulse, 25 ms) gave the highest survivorship, with a 1 dpf survival rate of 85% (Supp. Fig. S1). We then fixed the larvae from condition 3 and the NE control at the 3 dpf stage, and used confocal microscopy to confirm that the electroporation parameters from condition 3 had successfully delivered Dextran inside the embryos and that the fluorescence was distributed among most cells throughout development (Supp. Video S1-S2). The success of electroporation condition 3 in delivering Dextran gave us confidence to begin testing shRNA transfection in *H. symbiolongicarpus* embryos.

**Figure 2.**
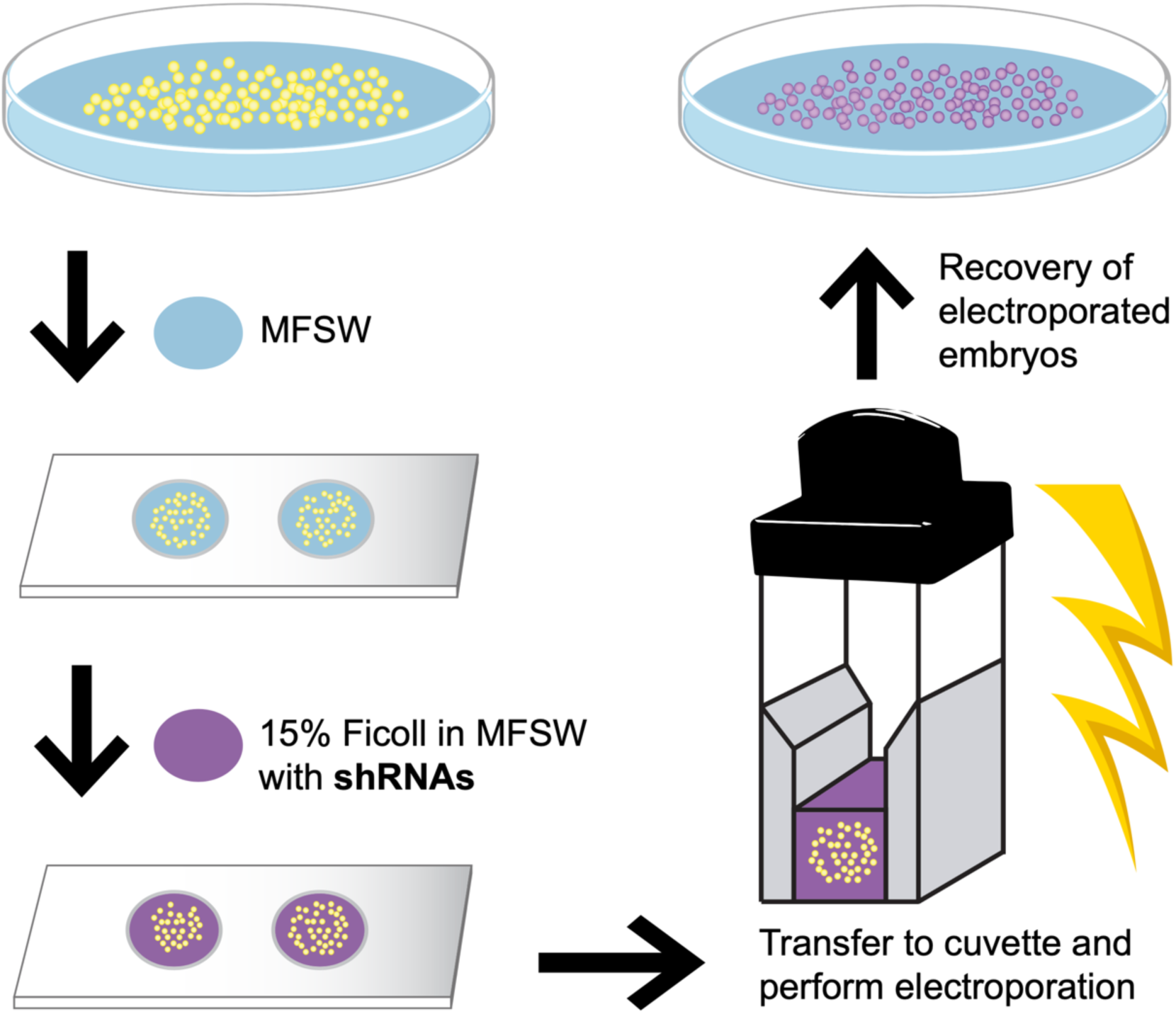
Overview of the shRNA electroporation procedure for *H. symbiolongicarpus* embryos. One-cell stage embryos are collected in a petri dish and transferred in a small volume of Millipore-filtered seawater (MFSW - blue) into wells of a depression slide. MFSW in each well is removed and replaced by 100μl of the electroporation solution, consisting of 15% Ficoll in MFSW containing the shRNAs (purple). The embryos in the electroporation solution are then transferred into an electroporation cuvette. The cuvette is placed inside the safety stand of the ECM 830 Square wave electroporation system (BTX) and electroporation of the embryos with the chosen parameters is carried out. Electroporated embryos are then carefully transferred to a petri dish with MFSW for their recovery and future phenotypic analyses. A detailed protocol can be found in Supp. File S1.

**Figure 3.**
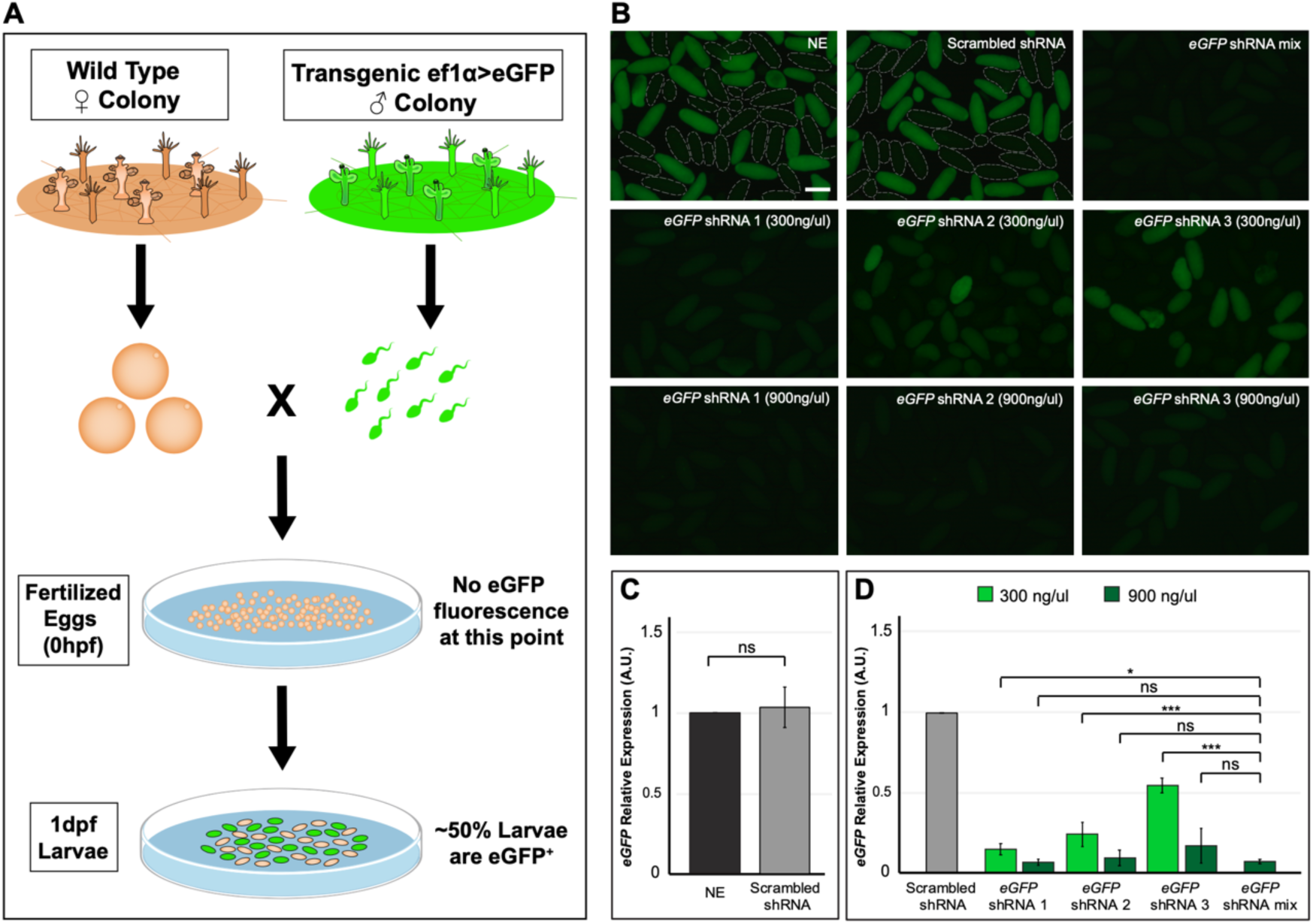
Visualization and quantification of *eGFP* knockdown. (**A**) Schematic depicting the cross performed leading to ∼50% of eGFP + larvae in the offspring due to Mendelian inheritance. This cross was carried out for all experiments involving the *eGFP* gene. (**B**) Representative fluorescence images of 1 dpf larvae for each of the nine experimental conditions shown. Note the reduction in eGFP fluorescence in the *eGFP* shRNA-electroporated conditions when compared to control larvae, with the *eGFP* shRNA mixture (mix) condition and the individual shRNAs at 900 ng/μl yielding the highest reductions. eGFP-larvae are outlined with dashed lines in the NE and scrambled shRNA controls for better clarity. NE = Non-Electroporated. (**C**) RT-qPCR results showing equivalent expression levels of *eGFP* mRNA in control samples. Scrambled shRNA *eGFP* expression levels were quantified relative to the NE control. (**D**) RT-qPCR results showing a clear reduction in the *eGFP* mRNA expression levels in all *eGFP* shRNA-electroporated conditions when compared to the scrambled shRNA control. When comparing the *eGFP* shRNA mixture condition with any of the three *eGFP* shRNAs electroporated separately at a concentration of 300 ng/μl each, the mixture yielded the highest *eGFP* transcript reduction of all, significantly lower than the rest. When comparing the *eGFP* shRNA mixture condition with any of the three *eGFP* shRNAs electroporated separately at a concentration of 900 ng/μl each, all conditions yielded similarly low levels of *eGFP* transcripts, indicating comparable knockdown efficiencies between these four conditions. For all *eGFP* shRNA experimental conditions, *eGFP* expression levels were quantified relative to the scrambled shRNA control. All RNA samples used for RT-qPCR analyses were extracted from 1 dpf larvae. Bar heights represent mean values of at least three independent experiments and error bars show standard errors of the mean. Significance levels: ∼∼∼ = p-value ≤ 0.01; ∼∼ = p-value ≤ 0.05; ∼ = p-value ≤ 0.1; ns = non-significant. Scale bar = 200μm.

**Figure 4.**
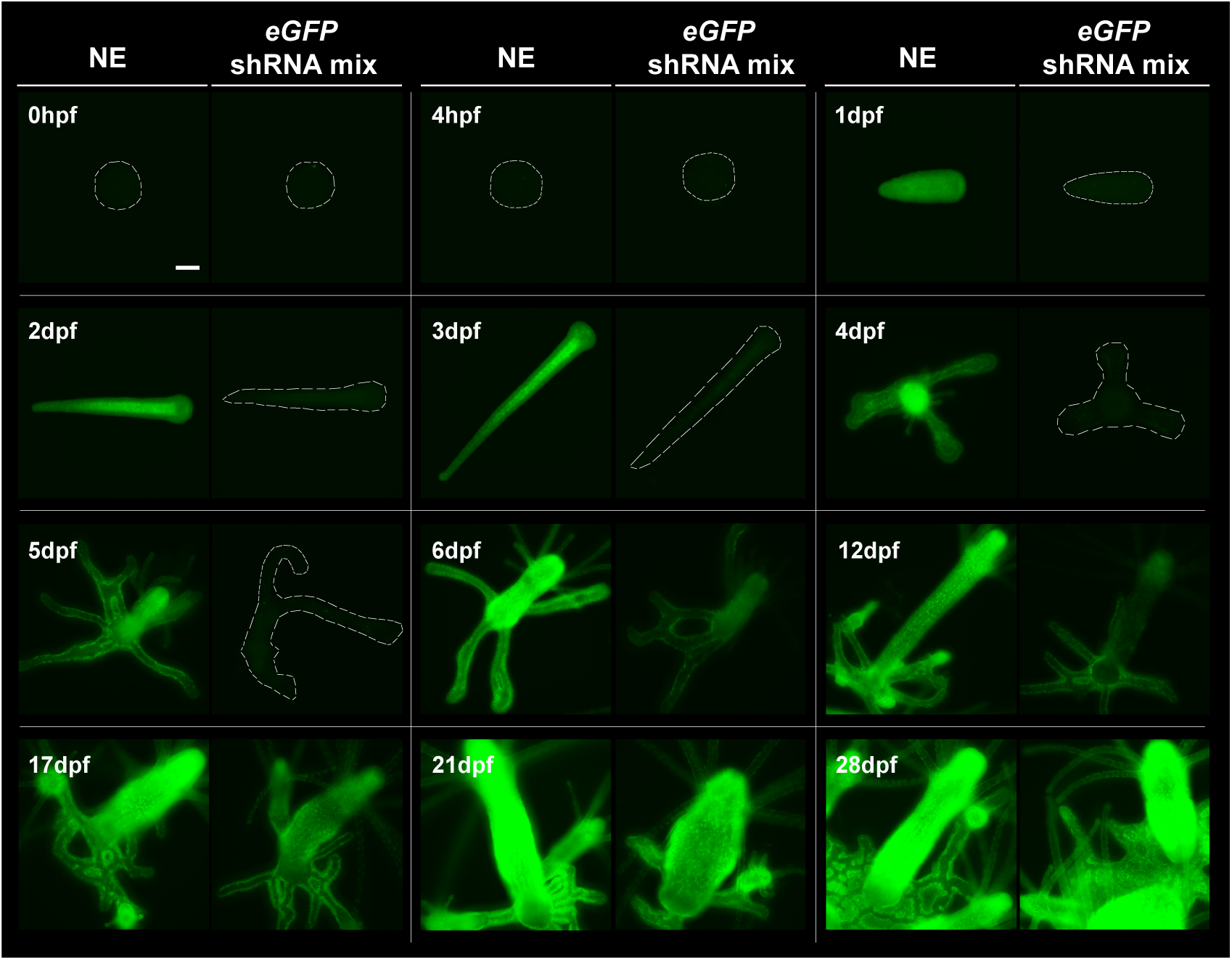
Fluorescence dynamics of *eGFP* knockdown over time. Representative fluorescence images from a larger number of samples (n > 30 in all cases) are shown for each selected timepoint and condition. ∼50% of NE control animals started appearing fluorescent by 1 dpf, and their fluorescence levels were saturated by 21 dpf. In contrast, the *eGFP* shRNA mixture-electroporated animals stayed nonfluorescent throughout embryonic development and during the first two days of primary polyp growth. At the 6 dpf timepoint, the fluorescence levels start to increase in ∼50% of the polyps, and it took up to 28 dpf for the fluorescence levels to be saturated. For clarity, embryos and polyps are outlined with dashed lines in all cases where eGFP fluorescence is not obvious. Scale bar = 100μm.

**Figure 5.**
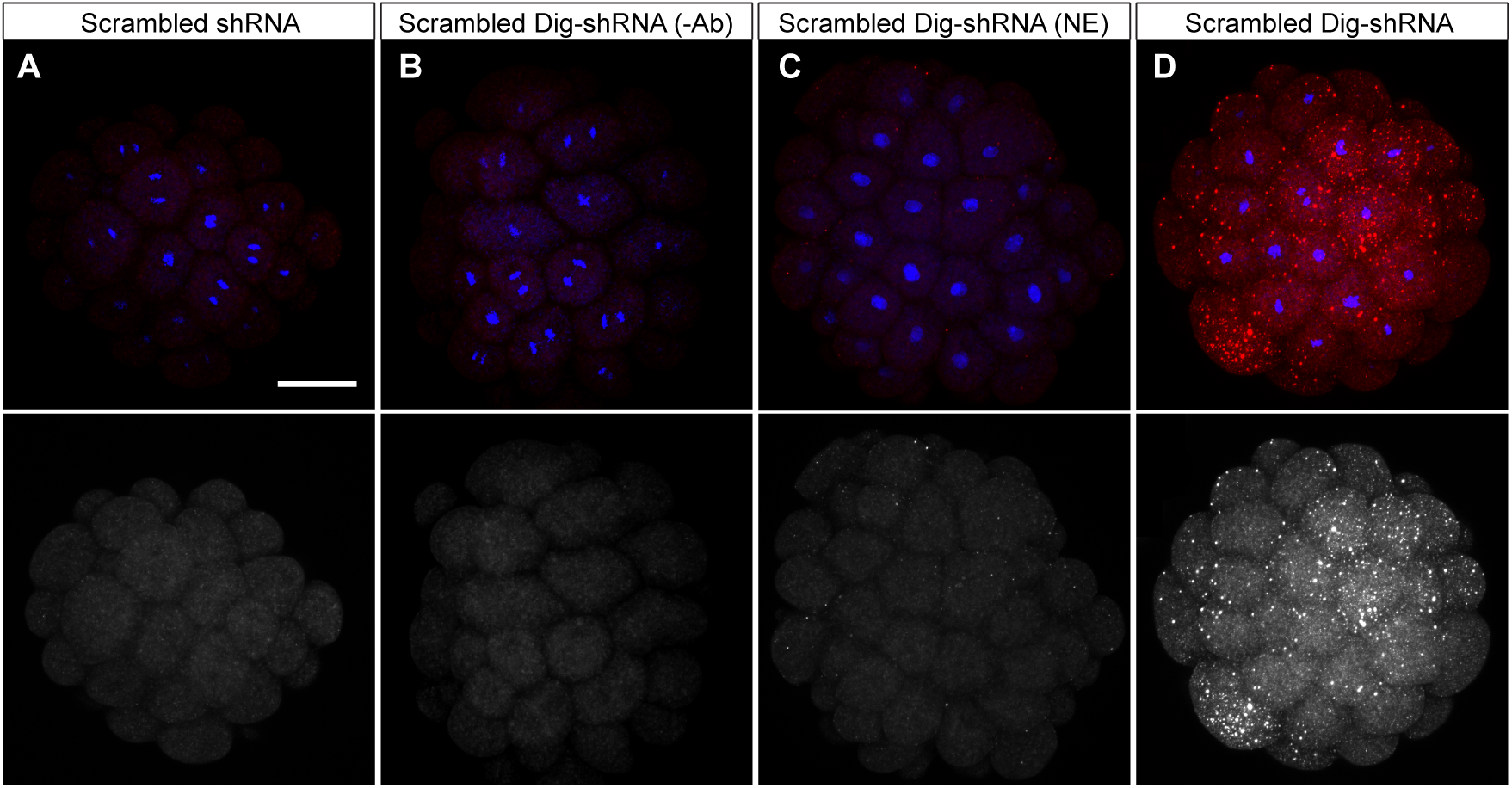
Visual confirmation of shRNA delivery into embryos via electroporation. Representative images of 4 hpf embryos are shown for each experimental condition. All images were projected from confocal z-stacks of ∼20μm. Hoechst staining of DNA is shown in blue (note some embryonic cells undergoing mitosis) and rhodamine-tyramide in red. The bottom row shows the single red channel in black and white for each condition to enhance contrast. In all cases, shRNAs were delivered at a concentration of 900 ng/μl. (**A**) Scrambled shRNA-electroporated embryo for which tyramide signal amplification (TSA) was carried out. (**B**) Digoxigenin-labeled scrambled shRNA-electroporated embryo for which TSA was performed but peroxidase-labeled anti-DIG antibody (Ab) was not added. Note the complete lack of rhodamine-tyramide signal in A-B controls. (**C**) Embryo that was soaked with digoxigenin-labeled scrambled shRNA for the length of an electroporation procedure (∼3min) but was not electroporated, and for which the TSA reaction was carried out. Note that some dots appear with the rhodamine-tyramide signal, mostly from shRNAs stuck on the surface but not inside the cells (see Supp. video S3). (**D**) Digoxigenin-labeled scrambled shRNA-electroporated embryo for which TSA was performed. Notice the intense fluorescence generated from the rhodamine-tyramide signal, largely coming from shRNAs that were delivered inside the one-cell stage embryonic stage via electroporation, and which stayed inside the cells throughout the first stages of cleavage (see Supp. video S4). Scale bar = 50μm.

**Figure 6.**
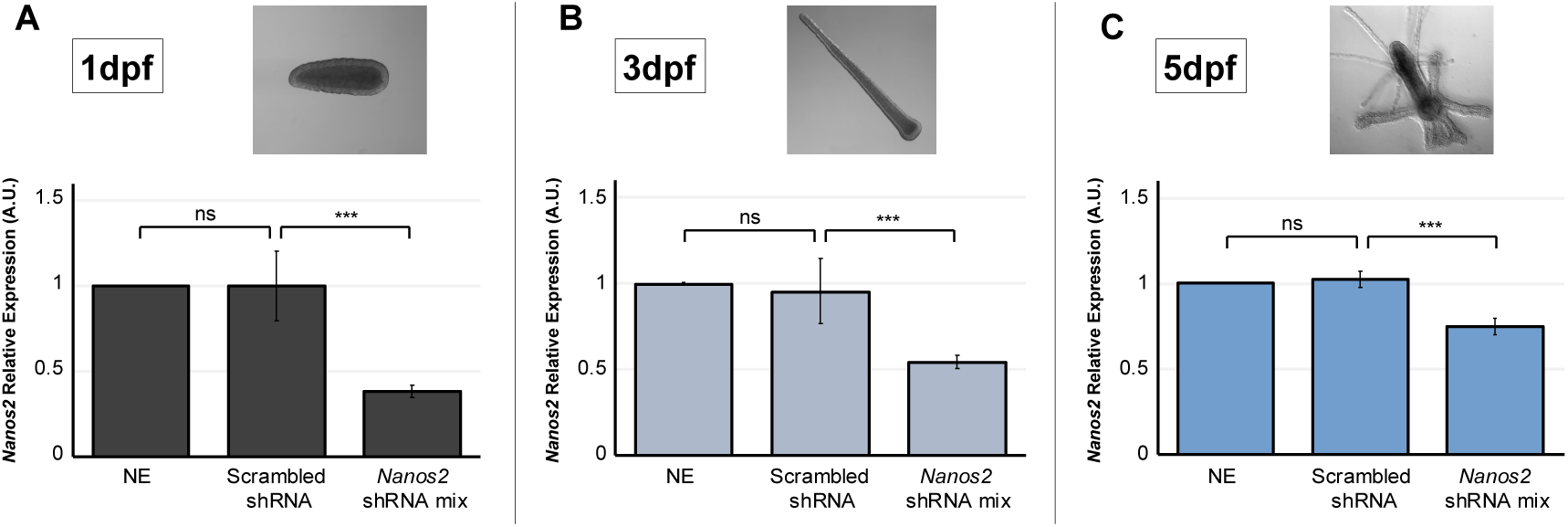
*Nanos2* knockdown quantification over time. RT-qPCR results from samples, showing comparable expression levels of *Nanos2* mRNA in both controls, but a significant decrease (p-value ≤0.01) of *Nanos2* mRNA expression levels when comparing *Nanos2* shRNA mixture-electroporated samples to the scrambled shRNA control at 1 dpf (**A**), 3 dpf (**B**), and 5 dpf (**C**). ns = non-significant. Images of how the animals looked at each RNA extraction timepoint are shown. In all cases, levels of *Nanos2* expression in scrambled shRNA and *Nanos2* shRNA mixture (mix) samples were quantified relative to the NE control. Bar heights represent mean values of three independent experiments and error bars show standard deviations.

**Figure 7.**
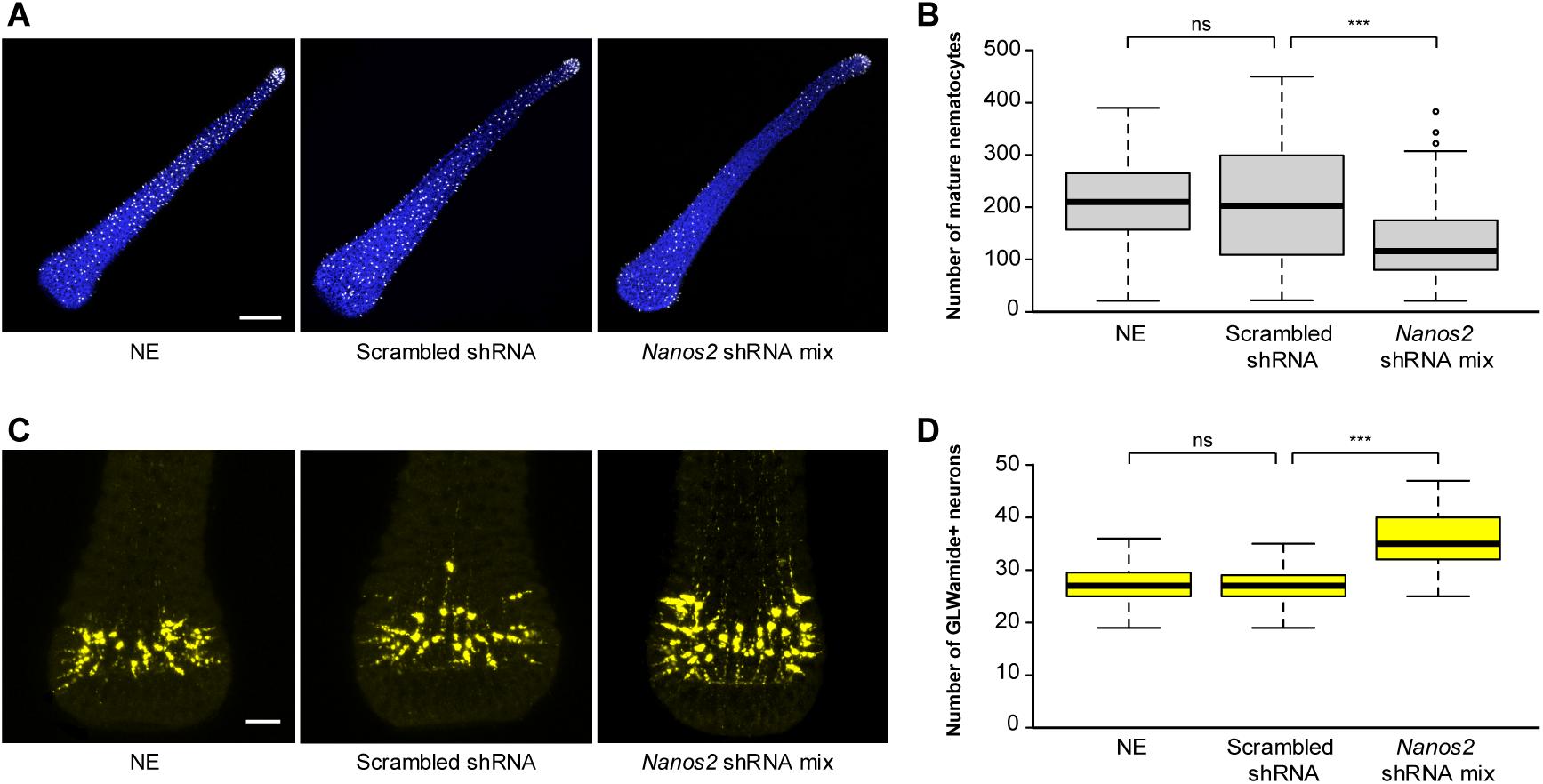
Phenotypic characterization of *Nanos2* knockdown larvae. (**A**) Representative images of 3 dpf larvae showing the mature nematocytes (white) on the larval surface for each of the three different conditions presented. Cell nuclei are stained in blue. Images were projected from confocal stacks of ∼10μm. (**B**) Box plot showing the number of mature nematocytes for each of the three conditions. Center lines show the medians; box limits indicate the 25th and 75th percentiles (first and third quartiles); whiskers extend 1.5 times the interquartile range from the 25th and 75th percentiles; outliers are represented by circles. For NE, n = 49; for scrambled shRNA, n = 53; for *Nanos2* shRNA mixture (mix), n = 65. (**C**) Representative images of 3 dpf larvae showing the GLWamide^+^ neurons (yellow) on the larval aboral region for each of the three different conditions. Images were projected from confocal stacks of ∼45μm. (**D**) Box plot showing the number of GLWamide^+^ neurons for each of the three conditions. Box plot characteristics as in (B). For NE, n = 72; for scrambled shRNA, n = 68; for *Nanos2* shRNA mixture, n = 75. Altogether, *Nanos2* knockdown yields a significant reduction (p-value ≤0.01) of mature nematocytes and a significant increase (p-value ≤0.01) of GLWamide^+^ neurons in the 3 dpf larvae. ns = non-significant. Scale bars = 100μm in (A) and 25μm in (C).

### Optimization of shRNA electroporation

Following our Dextran trials, we tested several electroporation conditions to assess the success in delivering shRNAs into one-cell stage embryos, and to determine the extent of shRNA-mediated gene knockdown in *H. symbiolongicarpus*. For these trials, we chose to target a gene with obvious phenotypic characteristics upon knockdown. We thus targeted the *eGFP* gene in the offspring of a cross between eggs produced by a wild type female and sperm from a transgenic *Eef1alpha>eGFP* male line (354-3)^10^.

The transgenic line was created via CRISPR/Cas9-mediated *eGFP* gene knockin to the endogenous housekeeping gene *Eef1alpha* (eukaryotic elongation factor 1 alpha) locus. This line was shown to express the exogenous *eGFP* gene in all somatic cells as well as in germ cells, thus allowing germline transmission of the transgene^10^. We consistently obtained approximately 50% of 1 dpf larvae that were eGFP^+^ from this wild type female by transgenic male cross (Fig. **3A**), as expected by Mendelian inheritance.

We designed three different shRNAs targeting various regions of the *eGFP* mRNA (see Methods). RNA-induced gene silencing in cnidarians requires nearly perfect complementarity^22,25^; therefore, off-targeting of shRNAs is highly unlikely. Nonetheless, we decided to exclude all sequences that had 16 complementary nucleotides or more with non-target genes to minimize the risk of off-target effects. Based on previous studies in the cnidarian *Nematostella vectensis*^22,23^, we determined that a concentration of 300 ng/μl for each shRNA should be effective in inducing the knockdown of most genes for several days in *H. symbiolongicarpus*. We decided to test a strategy of electroporating a mixture of three shRNAs at a time, each at a concentration of 300 ng/μl per shRNA (total concentration of 900 ng/µl), to yield high knockdown levels with very low risk of off-target effects. We reasoned that testing three shRNAs at once would be effective even if one shRNA was less active than the others or if the activity varied among shRNAs.

We carried out electroporation trials as detailed in **Figure 2** under varying electroporation parameters and experimental conditions (Supp. Fig. S2). *eGFP* shRNA mixtures were diluted in 15% Ficoll MFSW to a concentration of 300 ng/μl per shRNA (900 ng/μl total). In all cases, we assessed survivorship and eGFP fluorescence levels by counting the number of eGFP^+^ larvae at 1 dpf (Supp. Fig. S2). To evaluate any potential deleterious effects from high concentrations of shRNAs, we also tested two different control electroporation conditions where nuclease-free H_2_O was added instead of shRNAs (‘No shRNA’ controls). We observed that the survivorship levels in the ‘No shRNA’ controls were comparable to the same conditions where the *eGFP* shRNA mixture was added (Supp. Fig. S2), indicating that shRNA electroporation at a final concentration of 900 ng/μl was not toxic and did not affect survivorship levels of the animals. We also observed that electroporation of the *eGFP* shRNA mixture was successful in yielding lower levels of eGFP fluorescence than in controls in 1 dpf larvae under all conditions tested (Supp. Fig. S2). This result strongly suggested that *eGFP* knockdown could be accomplished through shRNA electroporation in *H. symbiolongicarpus* embryos. In agreement with the Dextran trials, the optimal electroporation condition for shRNA transfection, based on highest survivorship (∼85%) and lowest percentage of eGFP^+^ larvae (∼2% of ∼50% expected), was 25 V, 1 pulse, 25 ms (Condition A, Supp. Fig. S2). Thus, we used these electroporation parameters for all subsequent shRNA experiments.

We then tested whether electroporation of the *eGFP* shRNA mixture yielded a stronger reduction in eGFP levels than that of each of the three shRNAs (*eGFP* shRNAs 1-3) when electroporated separately. In one set of experiments, each shRNA was tested at the concentration they have in the mixture (i.e., 300 ng/μl each) and compared to the mixture (total concentration of 900 ng/µl). In another set of experiments, each shRNA was tested at the same final concentration as the mixture (i.e., 900 ng/µl each) and compared to the mixture (total concentration of 900 ng/µl). We included a NE control and a scrambled shRNA control for each experiment (Fig. **3B**). The scrambled shRNA consists of a random shRNA sequence that does not target any gene in the *Hydractinia* genome and serves as a negative shRNA control when added at the highest concentration of the experiment (900 ng/μl). We acquired fluorescent images of 1 dpf larvae for each of the experimental conditions. In all experiments, the NE and scrambled shRNA controls showed the expected percentage of highly fluorescent 1 dpf larvae (∼50%; Fig. **3B**). All electroporation conditions yielded a 75-97% survival rate at 1 dpf (Supp. Table S2). For the experiments where each individual shRNA was tested at a concentration of 300 ng/µl, *eGFP* shRNA 1 produced the most obvious reduction in eGFP fluorescence of the three individual shRNAs but the *eGFP* shRNA mixture appeared to show the strongest reduction of all conditions. For the experiments where each individual shRNA was tested at a concentration of 900 ng/µl, we observed that shRNAs 1 and 2 displayed virtually the same low level of fluorescence as the mixture, whereas embryos electroporated with shRNA 3 exhibited a slightly higher overall fluorescence level (Fig. **3B**).

To verify that the reduction in fluorescence levels was due to a reduction in *eGFP* transcript levels and to analyze the degree of gene-specific knockdown, we performed RT-qPCR analyses on 1 dpf larvae (Fig. **3C-D**). We first compared the *eGFP* mRNA levels between the NE and scrambled shRNA controls. As expected, the difference in the *eGFP* mRNA levels was not significant between the controls (Fig. **3C**). We then measured the *eGFP* mRNA levels for all experimental conditions relative to the scrambled shRNA control (Fig. **3D**). For all conditions, we found significant reductions in *eGFP* mRNA expression (Student’s T-test; p-value ≤ 0.01), with the *eGFP* shRNA mixture and shRNA 1 giving the most dramatic knockdowns in the set of experiments where each individual shRNA was tested at the lower 300 ng/µl concentration. When we compared each shRNA at 300 ng/µl concentration to the *eGFP* shRNA mixture, the mRNA levels in the mixture were significantly lower than in any of the individual *eGFP* shRNAs alone, although the difference between *eGFP* shRNA 1 and the mixture was less obvious (p-value ≤ 0.1), indicating that *eGFP* shRNA1 yielded the strongest knockdown of the three individual shRNAs (Fig. **3D**). In the set of experiments where the individual shRNAs were each at the higher concentration of 900 ng/µl, all shRNAs gave a knockdown similar to the mixture condition and the observed differences were not statistically significant (Fig **3D**). Thus, there did appear to be an effect of shRNA concentration on overall transcript levels, with the higher concentration of 900 ng/µl being more effective in reducing *eGFP* transcript levels for all individual shRNAs tested. Together, these results show that the reduction in eGFP fluorescence levels we observed following *eGFP* shRNA electroporation represent successful gene-specific knockdowns. The results also show that the *eGFP* shRNA mixture strategy produces a knockdown that is as effective and dramatic as the most efficient individual shRNA at the highest concentration. Given these results, we opted to use the shRNA mixture strategy for all subsequent experiments.

### Characterization of *eGFP* knockdown over time

To ensure that shRNA electroporation would not have harmful consequences on the developing animals in the long-term, we decided to follow embryos subjected to electroporation with the *eGFP* shRNA mixture through an extended time course. We compared a NE control sample to an *eGFP* shRNA mixture-electroporated sample for 28 dpf. We examined survivorship during the first three days of development in both samples and observed that, by 3 dpf, knockdown animals showed a 73.8% survival rate, as compared to a survival rate of 90.2% in the NE controls (Supp. Table S2). This 16.4% difference in survivorship between samples is likely explained by inherent electroporation-induced damage (Supp. Fig. S2). We determined that the highest embryo mortality occurred during the first two days of development in both NE and electroporated animals. The surviving knockdown larvae at each developmental stage did not exhibit any noticeable negative appearance when compared to the NE sample, and all surviving individuals became fully developed 3 dpf larvae (Fig. **4**). Both NE and 3 dpf knockdown larvae were competent to metamorphose into primary polyps upon CsCl stimulation (see Methods). The two samples had very similar metamorphosis rates (77.0% NE and 78.2% *eGFP* knockdown; Supp. Table S2), indicating no negative effect of shRNA electroporation on metamorphosis. Mortality after metamorphosis was virtually absent in both cases. Furthermore, all polyps from both samples were successfully fed with smashed brine shrimp and steadily grew to form polyp colonies (Fig. **4**). These results indicate that *eGFP* shRNA mixture-electroporated embryos are capable of fully developing into 3 dpf larvae, metamorphosing into primary polyps, feeding, and growing to eventually form a colony.

We were also interested in documenting the eGFP fluorescence dynamics over time in *eGFP* knockdown animals compared to the NE control, where the fluorescence level corresponds to the amount of eGFP protein inside the cells. To evaluate this, we obtained representative fluorescent images of both samples at different timepoints throughout the experiment and used fluorescence levels to understand the phenotype dynamics over time (Fig. **4**). We first fed the young primary polyps at 5 dpf, the stage by which their mouths are fully formed. Subsequent feedings were done every other day. We noticed that feeding had an obvious impact on polyp growth and apparently on increasing fluorescence levels when comparing samples at 5 dpf and 6 dpf (Fig. **4**).

While eGFP fluorescence was virtually absent in knockdown animals throughout development and in 4-5 dpf primary polyps, by 6 dpf we observed that fluorescence in the knockdown polyps started to return. The fluorescence levels kept increasing over time in both samples. eGFP fluorescence was saturated by 21 dpf in the NE control polyps, while it took up to 28 dpf for the *eGFP* shRNA mixture-electroporated samples to reach equivalent fluorescence levels (Fig. **4**). These results suggest that shRNA electroporation-induced knockdown animals need several weeks to fully recover eGFP protein levels from the targeted mRNA.

### Visual confirmation of successful shRNA transfection via electroporation

To have visual confirmation that electroporation was a successful method for shRNA delivery inside one-cell stage embryos of *H. symbiolongicarpus*, we synthesized scrambled shRNAs labeled with digoxigenin (Dig-shRNA) and electroporated embryos with this construct at 900 ng/μl in 15% Ficoll MFSW. Appropriate controls were performed (described in Fig. **5A-C** legend). By developing a tyramide signal amplification (TSA) reaction in fixed 4 hpf embryos (32-64 cell stages), we observed a strong fluorescence signal that indicated the presence of Dig-shRNA inside the embryo’s cells (Fig. **5D**; Supp. Video S3-S4). It appeared that every cell within each embryo that we observed showed fluorescence, although some cells displayed a stronger signal than others. This visual confirmation indicates that shRNAs were successfully delivered inside embryonic cells using our method.

### Endogenous gene knockdown through shRNA electroporation

To test whether our method could be used to knock down endogenous genes in *H. symbiolongicarpus*, we targeted the *Nanos2* gene, which is known to be essential for balancing the numbers of neurons and nematocytes (stinging cells) in *Hydractinia*^11^. *Nanos2* knockdown was previously achieved through microinjection of a translation blocking morpholino. We designed three different shRNAs targeting different regions of *Nanos2* mRNA and electroporated one cell-stage embryos with a *Nanos2* shRNA mixture (300 ng/μl per shRNA in 15% Ficoll MFSW, total concentration 900 ng/µl). We carried out RT-qPCR analyses using mRNA samples collected at different timepoints (1, 3, and 5 dpf) to evaluate *Nanos2* knockdown over time. In all cases, *Nanos2* shRNA mixture-electroporated samples showed a significant reduction in *Nanos2* mRNA levels when compared to scrambled shRNA controls (Fig. **6A-C**), indicating a successful knockdown of the gene. We observed that *Nanos2* transcript levels increased over time, with the lowest reduction in transcript detected at the latest timepoint we sampled, the post-metamorphic stage at 5 dpf (Fig. **6C**). Average survivorship in *Nanos2* knockdown 1 dpf larvae was ∼80% (Supp. Table S2) and all surviving larvae developed successfully and metamorphosed into polyps. We then performed phenotypic analyses in our knockdown animals by counting the number of mature nematocytes and GLWamide^+^ neurons in 3 dpf larvae, as well as the number of tentacles in 5 dpf primary polyps. We observed that *Nanos2* knockdown larvae displayed a significantly lower number of mature nematocytes, as well as a significantly higher number of GLWamide^+^ neurons than scrambled shRNA controls (Fig. **7A-D**). We also detected a significant decrease in tentacle numbers in *Nanos2* knockdown 5 dpf primary polyps compared to scrambled shRNA control polyps (Supp. Fig. S3). Additionally, we witnessed that most of the tentacles in the knockdown polyps were shorter than in the controls. Altogether, these results show that we successfully obtained the *Nanos2* knockdown phenotype previously described^11^ using our shRNA mixture electroporation strategy.

## Discussion

The ability to perform gene function analyses in a wide range of organisms across the animal tree is essential for understanding the biological functions of genes, as well as how these functions have changed throughout evolutionary time. In this study, we have determined the optimal electroporation parameters to successfully deliver shRNAs in embryos of the hydrozoan *H. symbiolongicarpus*: 25 V, 1 pulse, 25 ms length, 2mm-gap cuvettes, and 15% Ficoll in MFSW as the electroporation medium. Further, we have shown that our strategy represents a simple, fast, and robust method that is both scalable and efficient to temporarily disrupt mRNA function of both exogenous and endogenous genes in *H. symbiolongicarpus* embryos.

The method we present to induce gene knockdown in *H. symbiolongicarpus* embryos has several advantages, and electroporation represents an attractive alternative to other delivery methods such as microinjection or soaking. The design of shRNAs is straightforward (Supp. File S2) and their synthesis is inexpensive compared to other antisense molecules such as morpholinos^33^. Moreover, shRNAs are advantageous activators of the endogenous RNAi pathway since they retain a relatively low rate of degradation and turnover^17^. Another important benefit lies in the large number of embryos that can be efficiently transfected at a time via electroporation. We have found the trials with Dextran a particularly useful starting point for testing small-molecule delivery into *H. symbiolongicarpus* embryos and would encourage a similar approach for researchers developing electroporation of small molecules for their own research organisms of interest. Our strategy of labeling shRNAs with digoxigenin also proved to be a valuable approach, since it allowed us to obtain a visual confirmation of the successful shRNA transfection inside embryonic cells with the chosen electroporation parameters. We would like to highlight the importance of the addition of NE and scrambled shRNA controls in each experiment to determine the embryo damage caused by electroporation and the effect of shRNA transfection on embryo fitness, respectively. Using a mixture of three different shRNAs targeting different regions of the same gene has proven to be an effective strategy to induce knockdowns in *H. symbiolongicarpus* embryos. We found that testing one or several shRNAs individually at different concentrations is also a valid strategy, and that achieving the most efficient knockdown may require optimization of finding the best shRNA(s) and their optimal concentration for each target gene.

A 28-day time course experiment examined the long-term effects of shRNA electroporation on *H. symbiolongicarpus* as well as the eGFP fluorescence dynamics following knockdown of *eGFP* through time. We found that embryos subjected to shRNA electroporation could develop into larvae, undergo metamorphosis, and become healthy polyps capable of feeding and growing into young colonies, showing that shRNA electroporation does not produce long-term harmful effects in our animals. Feeding young polyps at day 5 enhanced polyp growth and fluorescence levels when comparing samples at 5 dpf (prior to feeding) and 6 dpf (one day after feeding; Fig. **4**), indicating that feeding had a clear effect on the metabolism of the polyps and on the synthesis of the eGFP protein. Fluorescence in the *eGFP* knockdown polyps started to return at 6 dpf, likely implying that the *eGFP* shRNAs inside the cells were starting to wear off or diminish their activity. Finally, we hypothesize that the difference in the timepoints in which NE control and *eGFP* knockdown colonies displayed saturated levels of fluorescence (21 and 28 dpf, respectively; Fig. **4**) could be explained by a delay in eGFP protein synthesis in the *eGFP* knockdown animals.

Our method has allowed us to recapitulate a morpholino microinjection-induced knockdown phenotype of the endogenous gene *Nanos2* that was previously shown in *Hydractinia echinata*^11^. In that study, the authors performed *in situ* hybridization targeting the mRNA from the neuropeptide precursor of RFamide to show the increase in the larvae neuron numbers upon *Nanos2* knockdown. Here, we carried out immunofluorescence analyses to detect GLWamide^+^ neurons, showing a significant increase of these neural cells in *Nanos2* knockdown larvae (Fig. **7**). Our results with a complementary strategy corroborate and enhance our knowledge of *Nanos2* being a key factor for the proper balancing of neurons and nematocytes in *Hydractinia*, since the mRNA disruption of *Nanos2* clearly increases the number of at least two different types of neuropeptidergic neurons, while reducing the number of mature nematocytes.

Based on our data regarding *Nanos2* and *eGFP* knockdowns, we estimate that our method can be used to assess the knockdown of genes throughout development, settlement, and metamorphosis, but it might present limitations starting at post-metamorphic stages. Metamorphosis represents a radical change in the life cycle of *Hydractinia*, encompassing cell rearrangements and apoptosis^34,35^ as well as high levels of cell proliferation^36^. Throughout this dramatic process, most transfected shRNAs are likely diluted and/or degraded, making it difficult to maintain their biological activity, and thus a strong knockdown effect, after metamorphosis takes place. Nevertheless, the length of a particular knockdown is presumably gene-dependent, and it may also depend on the efficiency of the initial mRNA disruption.

Studying a larger cross-section of research organisms strongly enables our understanding of biological processes in the broader context of evolution and contributes to our knowledge of basic biological mechanisms common to all animals^37,38^. With the advent of gene editing tools, the possibility of performing functional genomic analyses in a diversity of species has become remarkably accessible^39,40^. RNAi-based tools such as shRNAs, however, still remain a popular choice within the widening array of technologies available to disrupt gene function. Deciding which tool to choose for a particular experiment will depend on the biological question a researcher aims to answer, the gene targeted, and the life stage of the organism that one is targeting^41^.

Electroporation of shRNAs into unfertilized eggs has been recently described for the anthozoan cnidarian *Nematostella vectensis*^23^. The few differences between our method and the one presented for *N. vectensis*, such as electroporating fertilized eggs in *H. symbiolongicarpus* compared to unfertilized eggs in *N. vectensis*, are due to the distinct biology of the two species. *Hydractinia* diverged from *Nematostella* ∼600 million years ago^42^ so it is unsurprising that their biology is quite distinct. One key difference is that *Hydractinia* is a fully marine species while *Nematostella* is estuarine (cultured in 1/3 strength seawater), thus it cannot be taken for granted that shRNA electroporation will behave the same in these two distant cnidarian species. The community of cnidarian researchers is rapidly broadening and becoming more interdisciplinary^43^, highlighting the need to develop new methods to study gene function in multiple species. Our comprehensive testing and success in inducing gene-specific knockdowns via shRNA electroporation of fertilized eggs in *H. symbiolongicarpus*, therefore, represents a methodological springboard for researchers studying gene function throughout embryonic development, settlement, and metamorphosis in a variety of cnidarian species. We hope that our detailed methodology will inspire researchers who work with diverse research organisms, especially marine invertebrates, to use similar strategies to knock down genes of interest.

## Acknowledgements

We thank Steve Sanders and Matt Nicotra for providing the *H. symbiolongicarpus Eef1alpha>eGFP* (354-3) transgenic line, Mackenzie Simon-Collins for assistance with figures, Maddison Harman and Cassidy Manzonelli for *Hydractinia* animal husbandry, and Mark Martindale and members of the Martindale lab for support with reagents and equipment. We also thank Shuonan He and Ahmet Karabulut for helpful advice on shRNA and experimental design. Finally, we thank Andy Baxevanis for providing helpful comments on the manuscript. This work was funded by the NSF program “Enabling Discovery through Genomics tools – EDGE” (grant no. 1923259).

## Author contributions

G.Q-A. conceptualized and designed the project, performed all experiments, analyzed data, prepared figures, and wrote the manuscript. A.D. performed short hairpin RNA experiments and analyzed resulting data. K.L. performed dextran trials and initial short hairpin RNA experiments. J.W. assisted with short hairpin RNA experiments. C.E.S. conceptualized and designed the project and wrote the manuscript. All authors approved the final manuscript.

## Additional information

**Correspondence** and requests for materials should be addressed to G.Q-A. or C.E.S.

