## Supplementary material for "Gene knockdown via electroporation of short hairpin RNAs in embryos of the marine hydroid *Hydractinia symbiolongicarpus*": Supp File S1

**Protocol: shRNA electroporation in *Hydractinia* embryos**

**Equipment and Reagents**

Materials and reagents for embryo preparation:

- Artificial seawater (30ppt; Reef Crystals)
- Millipore-filtered artificial seawater (MFSW, 30ppt) -- Artificial seawater filtered with a Vacuum Filtration System (PES Membrane, 0.22um, 1000mL, Sterile; Argos Technologies)--
- Large plastic bins (3 liters, Tri State Plastics, Catalog No. 195-C)
- Cell strainer -size 70 μm- (white, sterile, individually wrapped; Corning)
- Glass bowls (Carolina, 4 1/2 in, 250 mL)
- Analog Orbital Shaker (Medium, 20 mm Amplitude, 115/230 VAC; Elmi Sky Line)
- Stereo Microscope (Stemi 508; Zeiss)
- Disposable Borosilicate Glass Pasteur Pipets -9 inches- (Fisherbrand)
- Micro Slides (Plain, Culture, Two Depression; Erie Scientific)
- Rain‑X® Original Glass Water Repellent (optional)

Materials and reagents for electroporation mixture preparation:

- Ficoll (PM 400, Type 400; Sigma-Aldrich)
- Millipore-filtered artificial seawater (MFSW, 30ppt)
- Conical Centrifuge Tubes (15ml; Falcon)
- LP Vortex Mixer (Continuous/Touch Mode, 0 to 3000 rpm; Thermo Fisher Scientific)
- Microcentrifuge tubes (1.5ml; Fisherbrand)
- Nuclease-Free water
- Purified shRNAs

Electroporation equipment:

- ECM 830 Square wave electroporation system and safety stand (Adjustable gap, 45-0207) (BTX)
- Electroporation Cuvettes (2mm, Blue Cap, Square Lid, Individually Wrapped, Sterile; Lee plastic company)
- Glass petri dishes (with lid, soda-lime glass, size 100 mm × 15 mm; BRAND)

**Tasks to do prior to an electroporation experiment**

- Make sure the temperature of the experimental lab is ~18ºC. If the temperature is higher the embryos will develop more quickly.
- Prepare all necessary equipment and reagents for the experiment in advance.
- Prepare 15% Ficoll in MFSW (“embryo suspension medium”):
  - In a 15ml conical/falcon tube, dissolve 0.75g Ficoll PM400 in 3.5ml MFSW. Vortex for 2 minutes and shake in a nutator for at least 30 minutes. Adjust the volume to 5ml by adding MFSW. Vortex for 1 more minute and let the tube settle at ~18ºC.

Larger volumes can be prepared, but we recommend 5ml final volume and to prepare fresh 15% Ficoll in MFSW every 2-3 weeks to avoid precipitation and contamination.

15% Ficoll in MFSW is a slightly viscous solution and will initially look cloudy, but it will gradually become transparent within ten minutes.

- Perform the calculations for the electroporation mixtures (including the shRNA dilutions) in advance. Here is an example of a calculation:

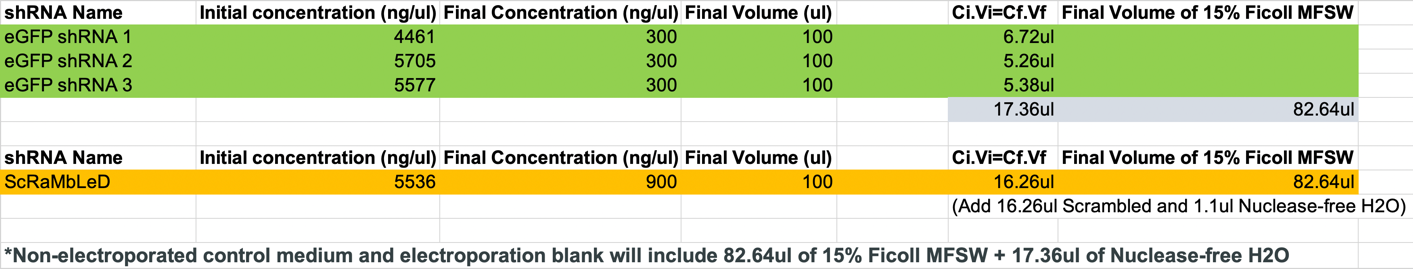

Note: A final volume of **100ul** of electroporation mixture has proven to be optimal for *Hydractinia* embryos successful electroporation in a 2mm cuvette.

Note: Always equilibrate salinity (e.g. Using the calculation example above, if you need ~17.4ul of *eGFP* shRNAs and only ~16.3ul of scrambled shRNA to reach the same final concentration, then add ~1.1ul more of Nuclease-free H2O in the scrambled tube to equilibrate final salinities).

- Turn on the ECM 830 system and set up the electroporation conditions to be used.

Optimized parameters for shRNA electroporation in *Hydractinia* embryos:

- Voltage (V) = 25V
- Number of pulses = 1
- Pulse length (ms) = 25ms
- Time span between pulses = NA
- Cuvette gap width (1, 2, 4mm) = 2mm

- Prepare the electroporation mixtures.
  - We recommend preparing all electroporation mixtures for a given experiment in 1.5ml sterile microcentrifuge tubes prior to the animals spawning. However, it is better to wait until spawning occurs to add the shRNAs into the electroporation mixtures to avoid wasting shRNAs in case the spawning is not good enough to carry out the experiment. This will also minimize the potential degradation of shRNAs in 15% Ficoll MFSW at 18ºC, since they will be added right before embryo electroporation.

Note: Collect the shRNAs from the -80C freezer and keep them on ice throughout the experiment. shRNAs are generally stable after 2-3 freeze-thaw cycles.

Note: Each experiment should include two types of negative controls, a scrambled shRNA electroporation control and a non-electroporated control. These controls will inform about the fertilization rate of that particular batch as well as of the embryo survival upon electroporation with shRNA that does not target any sequence in the animal’s genome.

- Optional: Use Rain-X® spray to coat the depression slides beforehand, swipe out the excess and let it dry for a few minutes. This helps to make a seawater drop that is easier to manipulate.

**Electroporation protocol**

1. Place *Hydractinia* female and male racks inside two separate large plastic bins containing artificial seawater (30ppt) while in the dark, then induce spawning by giving a direct light stimulus.
2. About 1h50min after the light stimulus, both females and males should have spawned eggs and sperm, respectively. Place the racks back into their culture tanks and keep the bins containing the male and female gametes in seawater.
3. Add seawater with sperm into the bin containing seawater with eggs.

Note: Amount of seawater with sperm added depends on the quantity of eggs spawned and the cloudiness of the sperm-containing seawater, which is also dependent on the spawning success.

4. Allow 15 minutes for fertilization to occur.

Note: During this time, add the shRNAs into the appropriate electroporation mixtures and mix by pipetting up and down until solution is homogeneous. Avoid creating bubbles.

Note: The ECM830 system needs a ‘Blank’ step before running a program. Therefore, an extra cuvette with the same amount and proportions of 15% Ficoll MFSW / Nuclease-free water is needed in parallel with your sample for each different electroporation parameters chosen. If all of your samples are using the same electroporation parameters, you will only need to run the Blank step once. We recommend to prepare and run the ‘Blank’ for the chosen electroporation condition during these 15 minutes.

5. Collect the fertilized eggs (one-cell stage embryos) by pouring the seawater through a sterile cell strainer (70um mesh) to catch the eggs. Recover fertilized eggs in a glass bowl filled to 1/3^rd^ with seawater and leave it on an orbital shaker (~90rpm) for ~1 minute. Altogether, this process takes ~5 minutes.
6. Using a Stereo Microscope, collect the fertilized eggs (up to 800 per experimental condition) with a glass Pasteur pipette and place them in different wells of glass depression slides.
7. Focusing on one condition at a time, get rid of most leftover seawater in the well to minimize the dilution of your electroporation mixtures, but never let the embryos dry out. Immediately after removing seawater, add the respective electroporation mixture into the well containing the fertilized eggs and delicately flush the 100ul drop with a glass Pasteur pipette to mix and homogenize.

Note: The fertilized eggs should float in the electroporation mixture thanks to the high concentration of Ficoll. If they do not, prepare fresh 15% Ficoll MFSW. The solution should look transparent and clean.

8. Carefully transfer the 100ul drop of electroporation mixture containing the fertilized eggs into an electroporation cuvette (2mm gap) with a glass Pasteur pipette.

Note: Use a black background to see the embryos being transferred into the cuvette.

Note: Avoid creating bubbles.

Note: Avoid touching the walls of the cuvette with the pipette. If fertilized eggs get stuck on the sides of the cuvette chamber they will not be properly electroporated.

9. Delicately flick the cuvette a few times to evenly distribute the fertilized eggs floating inside. Right after this, place the cuvette inside the safety stand connected to the ECM 830 electroporator.
10. Perform the electroporation with the previously chosen parameters by clicking ‘GO’ on the screen.
11. After electroporation is complete, remove the cuvette from the safety stand. Small bubbles should be seen on the walls of the cuvette chamber.
12. Very gently, transfer the electroporated embryos with a glass Pasteur pipette from the cuvette to a large glass petri dish filled with MFSW.

Note: It is crucial to do this step carefully and to evenly distribute the electroporated embryos in the big glass petri dish for them to develop properly. It is useful to pipette a small amount of MFSW from petri dish into the cuvette to gently flush the embryos, collect a small number of embryos, and then gently place in petri dish. Repeat 2-4 times to gently collect all embryos.

Note: For this step it may be useful to have a second person perform to be able to carry out electroporations for several conditions prior to the embryos cleaving.

13. Repeat steps 7-12 for each sample and chosen experimental condition.

Note: *Hydractinia* embryos take ~50 minutes to start cleaving when kept at 18ºC, normally allowing time to perform electroporations on up to 6 different samples per experiment.

Note: The electroporation cuvettes can be washed with ultrapure H_2_O and EtOH 100%, let dry for at least 48 hours and be reused again for up to 10 times.

14. Allow the embryos to recover and develop in the large glass petri dishes for several hours without disturbing them.
15. By the end of the day, transfer all healthy, developing embryos to a new glass petri dish. This will ensure an overall better development of the surviving embryos.

Note: Repeat the same procedure of transferring healthy, developing embryos 24 hours after electroporation to ensure proper development of the surviving embryos.

16. Perform detailed phenotypic studies at different timepoints depending on the biological question to be answered.
