## Supplementary material for "Gene knockdown via electroporation of short hairpin RNAs in embryos of the marine hydroid *Hydractinia symbiolongicarpus*": Supp File S2

**shRNA Design**

The first step is to find siRNAs targeting your sequence of interest (e.g. *eGFP*). For this step, we recommend the siRNA Wizard web interface from *Invivogen*: <http://www.invivogen.com/sirnawizard/design.php>

- Set motif size to 19 nucleotides.
- Paste the cDNA sequence of your gene of interest (e.g. *eGFP*) and click search.
- The browser will give you all the potential forward strand sequences targeting your gene of interest, the coordinates of where they are located inside your cDNA sequence, and the %GC content of each of them.

**eGFP**

Top of Form

| Sequence | Start | GC% | Design | Scrambled |
| --- | --- | --- | --- | --- |
| GAATTAGATGGTGATGTTA | 49 | 31.58 |  |  |
| **GTGAAGGTGATGCAACATA** | 98 | 42.11 |  |  |
| **GCCAACACTTGTCACTACT** | 171 | 47.37 |  |  |
| GGTTATGTACAGGAAAGAA | 271 | 36.84 |  |  |
| GGTGATACCCTTGTTAATA | 346 | 36.84 |  |  |
| GGAATACAACTATAACTCA | 423 | 31.58 |  |  |
| **GCGTTCAACTAGCAGACCA** | 524 | 52.63 |  |  |
| GCAGACCATTATCAACAAA | 535 | 36.84 |  |  |
| GGCATGGATGAACTATACA | 694 | 42.11 |  |  |
| GCATGGATGAACTATACAA | 695 | 36.84 |  |  |

Bottom of Form

Select three forward strand sequences targeting your gene of interest (our selections are shown here in bold) based on the following criteria:

- Overall GC content between 30-55% (based on the siRNA Wizard web interface from *Invivogen,* sequences with lower GC content seem to be more active than those with higher GC content)
- Low GC content on its 3’ end (helps RISC complex to load the reverse strand instead of the forward strand)
- Avoid sequences with regions of low complexity
- Target genes in the FIRST half of their sequence if possible (to avoid any potential translation of biologically active protein domains even if the mRNA is cleaved by the RISC complex)
  - NOTE: for qPCR primers, target the SECOND half of the sequence, or at least downstream from where the shRNAs are designed.

Next, BLAST the selected forward strands against your species’ genome gene models and Transcriptome. **To minimize potential off-targets, filter out all sequences that have 16 complementary nucleotides or more with non-target genes**.

If available, also BLAST the selected forward strands against your species’ miRNA database. If any of your chosen forward strands yields a match to a miRNA, discard it and select a new one.

Once the 3 best forward strands have been selected, use software such as Geneious® to create the template of your shRNAs:

A) Generate the reverse strand (the reverse complement of your forward strand)
B) Add the hairpin loop linker in between the forward and reverse strands (in our study, we have used the same linker that was employed in the shRNA design to target genes of the cnidarian *Nematostella vectensis*^1,2^)
C) Include two additional thymidines (TT) at the 3’ end of the template to mimic the endogenous pre-miRNA structure. The total length of the template should be now **49 nucleotides**.

- Here is an example of one shRNA template targeting the *eGFP* gene:

**5’ – GTGAAGGTGATGCAACATATTCAAGAGATATGTTGCATCACCTTCACTT – 3’**

Then, to predict whether your shRNA is going to properly fold *in vitro*, we recommend predicting its secondary structure by using the RNA mfold web server: <http://unafold.rna.albany.edu/?q=mfold>. Click on the ‘RNA folding form’ and paste the complete sequence. Perform prediction keeping all settings as default, click ‘Fold RNA’. Click on ‘Structure 1’ to see the fold.

The RNA mfold web server should predict the formation of a highly stable hairpin structure. If not, discard the sequence and select a new one.

Finally, include the T7 RNA polymerase promoter at the 5’ of your shRNA template. The total length of the template should be now **66 nucleotides**.

- Here is an example of a DNA template for *in vitro* transcription of an shRNA targeting the *eGFP* gene:

**5’ – TAATACGACTCACTATAGTGAAGGTGATGCAACATATTCAAGAGATATGTTGCATCACCTTCACTT – 3’**

A dsDNA template is needed for the *in vitro* transcription of shRNAs. Thus, a reverse complement of the 66 nucleotide-long sequences also needs to be designed. Both forward and reverse ssDNA oligonucleotides can then be ordered.

**Examples**

**Construct *eGFP* shRNA #1**       siRNA GC%: 42.11      Position: 98

Hairpin structure

5' GTGAAGGTGATGCAACATATTCAAGAGATATGTTGCATCACCTTCACTT 3'

- No hits found either in the Transcriptome or in the Genome of *H.symbiolongicarpus* that have complementarity of 16 nucleotides or more to the forward strand
- No hits found in the *H.symbiolongicarpus* miRNA database

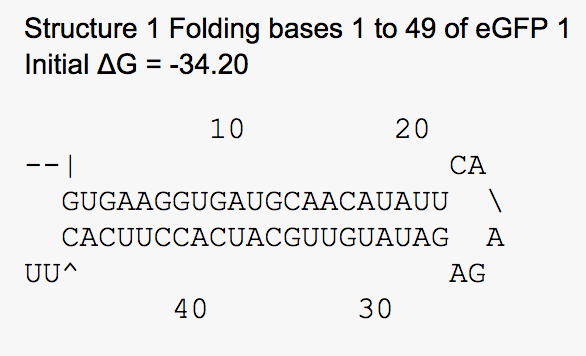

Oligo eGFP #1 (For)

5' TAATACGACTCACTATAGTGAAGGTGATGCAACATATTCAAGAGATATGTTGCATCACCTTCACTT 3'

Oligo eGFP #1 (Rev)

5' AAGTGAAGGTGATGCAACATATCTCTTGAATATGTTGCATCACCTTCACTATAGTGAGTCGTATTA 3'

**Construct *eGFP* shRNA #2**       siRNA GC%: 47.37      Position: 171

Hairpin structure

5' GCCAACACTTGTCACTACTTTCAAGAGAAGTAGTGACAAGTGTTGGCTT 3'

- No hits found either in the Transcriptome or in the Genome of *H.symbiolongicarpus* that have complementarity of 16 nucleotides or more to the forward strand
- No hits found in the *H.symbiolongicarpus* miRNA database

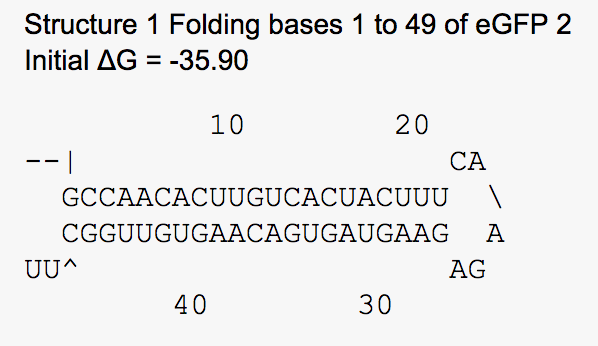

Oligo eGFP #2 (For)

5' TAATACGACTCACTATAGCCAACACTTGTCACTACTTTCAAGAGAAGTAGTGACAAGTGTTGGCTT 3'

Oligo eGFP #2 (Rev)

5' AAGCCAACACTTGTCACTACTTCTCTTGAAAGTAGTGACAAGTGTTGGCTATAGTGAGTCGTATTA 3'

**Construct *eGFP* shRNA #3**      siRNA GC%: 52.63      Position: 524

Hairpin structure

5' GCGTTCAACTAGCAGACCATTCAAGAGATGGTCTGCTAGTTGAACGCTT 3'

- No hits found either in the Transcriptome or in the Genome of *H.symbiolongicarpus* that have complementarity of 16 nucleotides or more to the forward strand
- No hits found in the *H.symbiolongicarpus* miRNA database

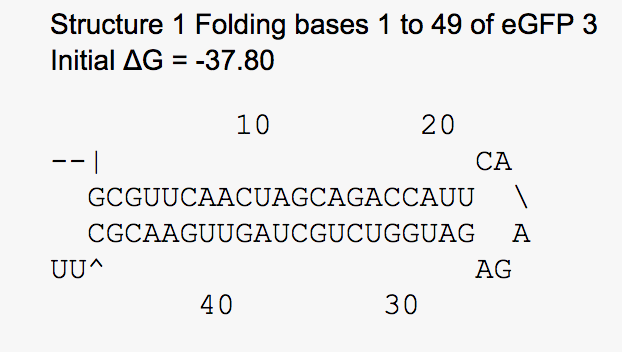

Oligo eGFP #3 (For)

5' TAATACGACTCACTATAGCGTTCAACTAGCAGACCATTCAAGAGATGGTCTGCTAGTTGAACGCTT 3'

Oligo eGFP #3 (Rev)

5' AAGCGTTCAACTAGCAGACCATCTCTTGAATGGTCTGCTAGTTGAACGCTATAGTGAGTCGTATTA 3'
