## Supplementary Information for "Gene knockdown via electroporation of short hairpin RNAs in embryos of the marine hydroid *Hydractinia symbiolongicarpus*"

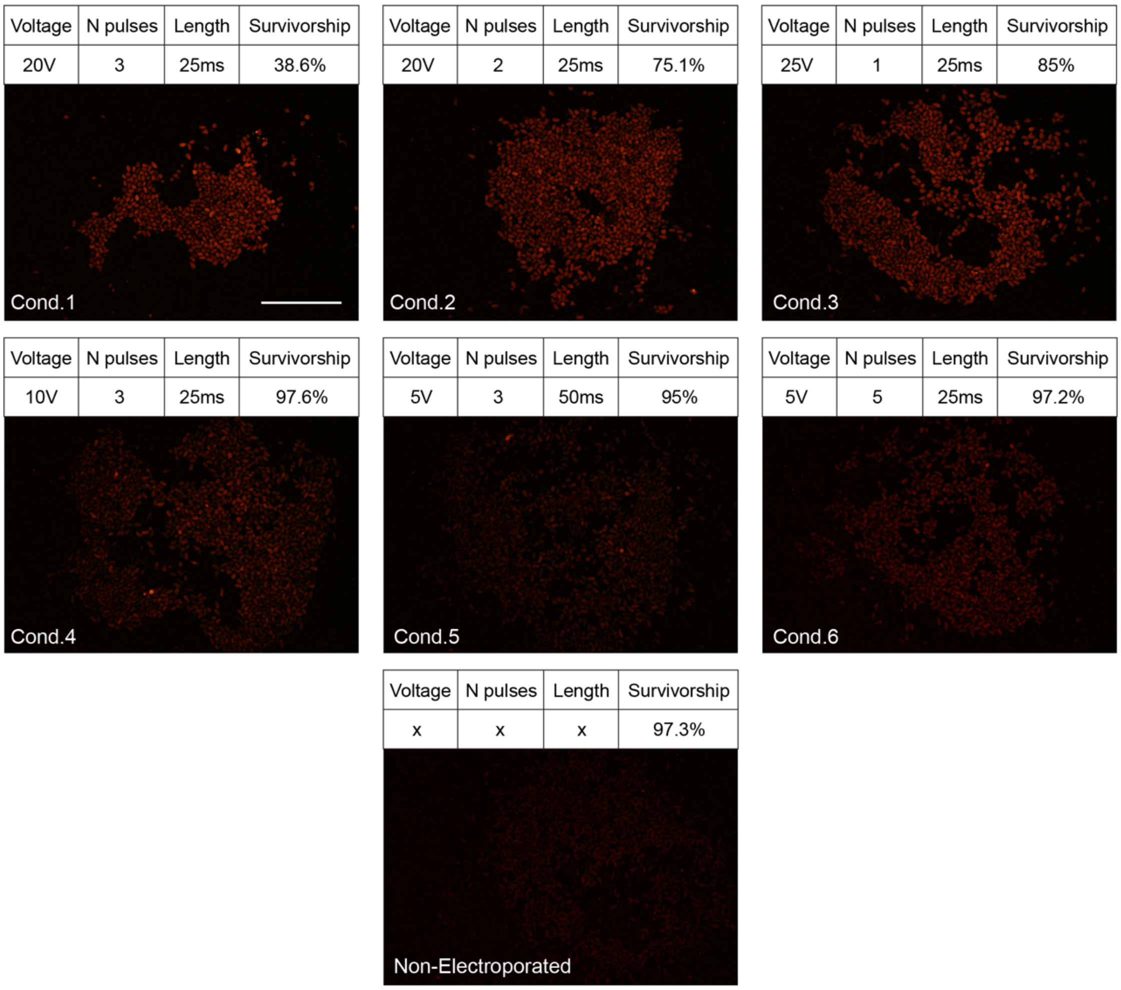


Supplementary Figure S1. **Dextran embryo transfection under different electroporation conditions**. Conditions (cond.) 1-6 account for different electroporation parameters (voltage, number of pulses, and pulse length) tested to deliver Dextran inside one-cell stage embryos. Images show 1 dpf larvae displaying different levels of Dextran fluorescence (red), which depended on how successful the delivery was from each condition. Note the almost complete lack of fluorescence in the non-electroporated control, indicating a lack of Dextran delivery inside the embryos. Survivorship at 1 dpf for each condition is also shown. Based on the fluorescence levels and the survival rate, condition 3 gave the most successful electroporation parameters for Dextran delivery into *H. symbiolongicarpus* embryos. Scale bar = 500µm.

**A**

| | Non-elec.1 | Non-elec.2 | No shRNA Ctrl (A) | No shRNA Ctrl (B) | A.1 | A.2 | B | C | D | E | F | G | H |
|---|---|---|---|---|---|---|---|---|---|---|---|---|---|
| Voltage (V) | N/A | N/A | 25 | 20 | 25 | 25 | 20 | 20 | 18 | 23 | 20 | 15 | 10 |
| #pulses | N/A | N/A | 1 | 3 | 1 | 1 | 3 | 2 | 2 | 1 | 1 | 3 | 6 |
| Pulse duration (ms) | N/A | N/A | 25 | 25 | 25 | 25 | 25 | 25 | 25 | 25 | 25 | 25 | 25 |

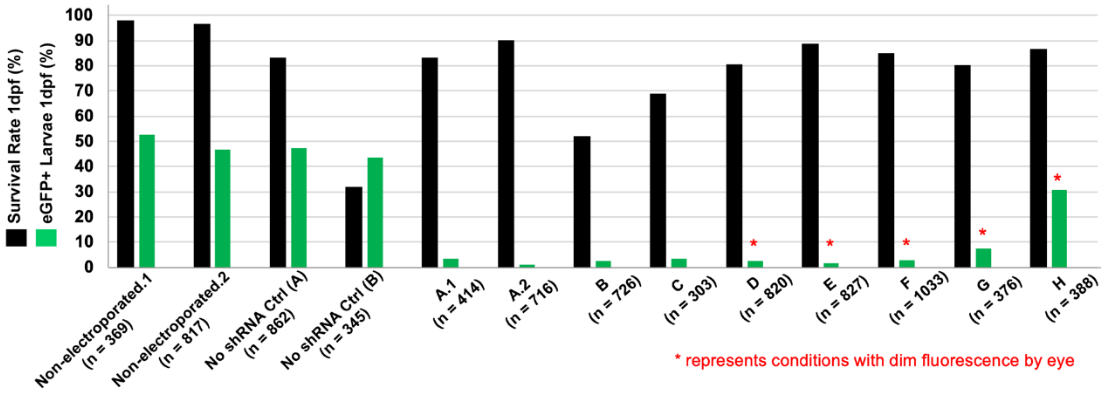


**B**

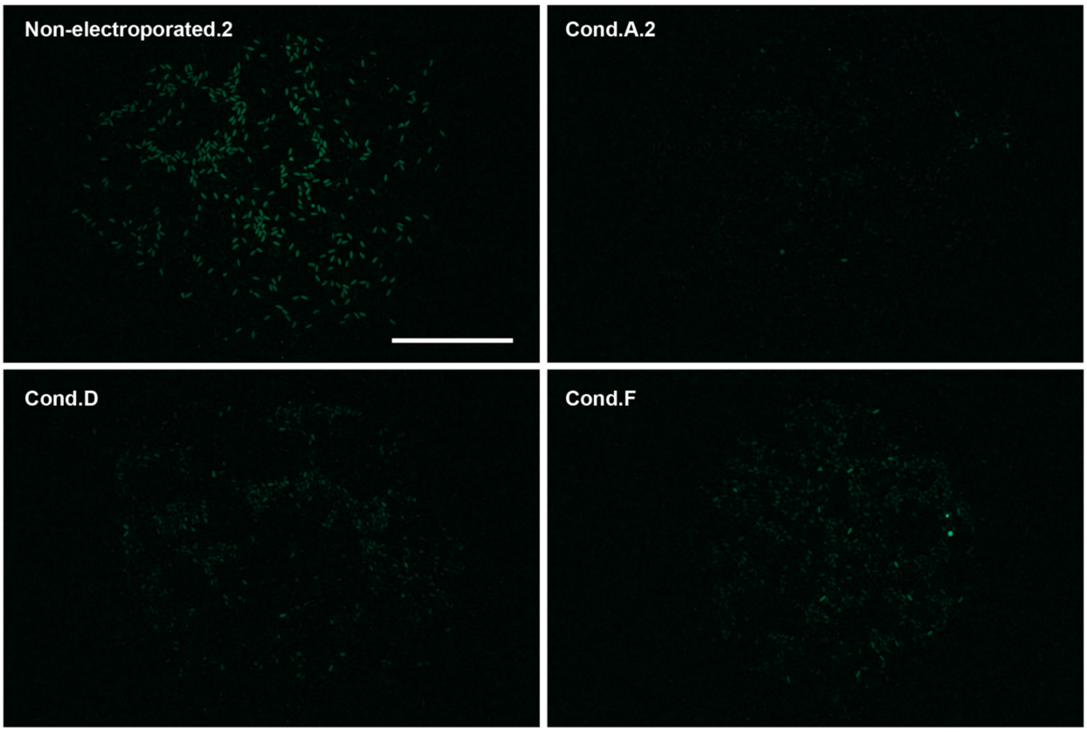


Supplementary Figure S2. **Survivorship and eGFP$^{+}$ larvae percentage for different electroporation conditions**. (**A**) Double-bar graph showing the survival rate and percentage of eGFP$^{+}$ larvae at 1 dpf after different electroporation conditions were used on fertilized eggs. The table above the graph containing the names of each condition and the electroporation parameters used corresponds vertically with the X axis of the graph. The names of the conditions that were repeated more than once are followed by a dot and a number, indicating the number of experiments where they were used. The number of electroporated embryos is shown for each condition at the bottom of the graph. Conditions A-H contained *eGFP* shRNA mixture in the electroporation solution. Note how the non-electroporated and the no shRNA controls present ~50% $eGFP^+$ larvae by 1 dpf, as expected by Mendelian inheritance, whereas conditions A-H always show lower percentages. Conditions presenting dim fluorescence seen by eye (D-H) are marked with a red asterisk and represent those for which dimly fluorescent larvae could be observed under a stereoscope, although this dim fluorescence could not be captured by the ImageJ software following our counting protocol (see Methods). (**B**) Fluorescence images of 1 dpf larvae for the conditions (cond.) presented. Notice the dimly fluorescent larvae in conditions D and F, likely indicating a less efficient delivery of *eGFP* shRNA mixture inside the embryos than in condition A. Based on the high survivorship and the low percentage of $eGFP^+$ larvae observed, condition A gave the most successful electroporation parameters for shRNA delivery into *H. symbiolongicarpus* embryos. Scale bar = 500μm.

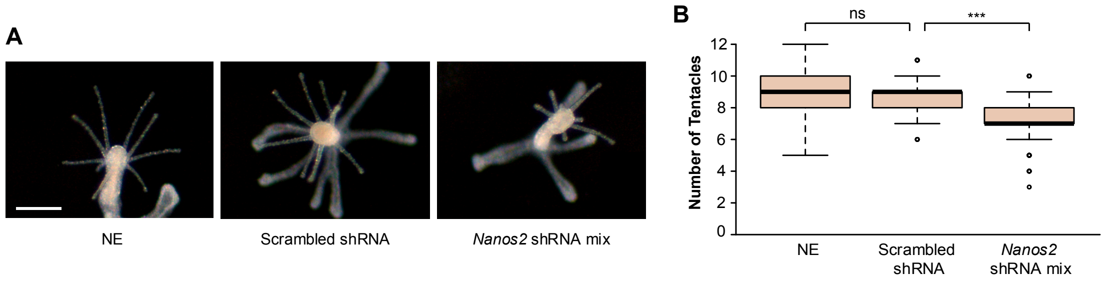


Supplementary Figure S3. **Tentacle number reduction in *Nanos2* knockdown animals**. (**A**) Representative images of 5 dpf primary polyps (2 days post-metamorphosis) for each of the three different conditions described. (**B**) Box plot showing the number of tentacles for each of the three displayed conditions. Center lines show the medians; box limits indicate the 25th and 75th percentiles (first and third quartiles); whiskers extend 1.5 times the interquartile range from the 25th and 75th percentiles; outliers are represented by circles. For NE, n = 201; for scrambled shRNA, n = 200; for *Nanos2* shRNA mixture (mix), n = 200. *Nanos2* knockdown yields a significant reduction (p-value ≤0.01) in the number of primary polyp tentacles. ns = non-significant. Scale bar = 250μm.

Supplementary Video S1. Representative 3 dpf larva from a non-electroporated control where one-cell stage embryos were soaked in 1mg/ml Dextran in 15% Ficoll MFSW for ~3min, the approximate length of an electroporation procedure. The video shows a sequence of images taken from the surface of the larva to a mid-region where the endoderm is visible. This movie was generated at a frame rate of 2 fps in ImageJ from a 20µm confocal z-stack. Nuclei (blue), F-actin (green), Dextran (red).

Supplementary Video S2. Representative 3 dpf larva from an experiment where one-cell stage embryos were electroporated with 1mg/ml Dextran in 15% Ficoll MFSW. The video shows a sequence of images taken from the surface of the larva to a mid-region where the endoderm is visible. This movie was generated at a frame rate of 2 fps in ImageJ from a 30µm confocal z-stack. Nuclei (blue), F-actin (green), Dextran (red).

Supplementary Video S3. Representative 4 hpf embryo from a non-electroporated control where one-cell stage embryos were soaked in 900 ng/µl Digoxigenin-labeled scrambled shRNA in 15% Ficoll MFSW for ~3min, the approximate length of an electroporation procedure. Prior to imaging, the 4 hpf embryos were fixed, then incubated with peroxidase-labeled anti-DIG antibody, then tyramide signal amplification with rhodamine-tyramide was carried out (see Methods). The video shows a sequence of images taken from the surface of the embryo to a mid-region. This movie was generated at a frame rate of 2 fps in ImageJ from a 15µm confocal z-stack. Nuclei (blue), rhodamine-tyramide (red).

Supplementary Video S4. Representative 4 hpf embryo from an experiment where one-cell stage embryos were electroporated with 900 ng/µl Digoxigenin-labeled scrambled shRNA in 15% Ficoll MFSW. Prior to imaging, the 4 hpf embryos were fixed, then incubated with peroxidase-labeled anti-DIG antibody, then tyramide signal amplification with rhodamine-tyramide was carried out (see Methods). The video shows a sequence of images taken from the surface of the embryo until to a mid-region. This movie was generated at a frame rate of 2 fps in ImageJ from a 25µm confocal z-stack. Nuclei (blue), rhodamine-tyramide (red).

Supplementary File S1. Detailed protocol for shRNA electroporation in *H. symbiolongicarpus* embryos.

Supplementary File S2. Detailed protocol for shRNA design.

Supplementary Table S1. List of the shRNA oligonucleotide sequences used for *in vitro* transcription of shRNAs and of the RT-qPCR primer sequences used in this study.

Supplementary Table S2. Embryo survival rate for different shRNA electroporation experiments targeting different genes.
